# PsiCLIP reveals dynamic RNA binding by DEAH-box helicases before and after exon ligation

**DOI:** 10.1101/2020.03.15.992701

**Authors:** Lisa M. Strittmatter, Charlotte Capitanchik, Andrew J. Newman, Martina Hallegger, Christine M. Norman, Sebastian M. Fica, Chris Oubridge, Nicholas M. Luscombe, Jernej Ule, Kiyoshi Nagai

**Author notes:** These authors contributed equally (Lisa M. Strittmatter, Charlotte Capitanchik). Joint corresponding authors. The authors dedicate this article to the memory of Kiyoshi Nagai (1949–2019).

## Abstract

Eight RNA helicases remodel the spliceosome to effect pre-mRNA splicing but their mechanism of action remains poorly understood. We have developed “purified spliceosome iCLIP” (psiCLIP) to define helicase-RNA contacts in specific spliceosomal states. psiCLIP reveals previously unappreciated dynamics of spliceosomal helicases. The binding profile of the helicase Prp16 is influenced by the distance between the branch-point and 3’ splice site, while Prp22 binds diffusely on the intron before exon ligation but switches to more narrow binding downstream of the exon junction after exon ligation. Notably, depletion of the exon-ligation factor Prp18 destabilizes Prp22 binding to the pre-mRNA, demonstrating that psiCLIP can be used to study the relationships between helicases and auxiliary splicing factors. Thus, psiCLIP is sensitive to spliceosome dynamics and complements the insights from structural and imaging studies by providing crucial positional information on helicase-RNA contacts during spliceosomal remodeling.

## Introduction

Splicing is an essential step of pre-mRNA processing in eukaryotes, in which non-coding introns are removed and coding exons are ligated. This reaction is catalyzed by a large, dynamic molecular machine called the spliceosome, which comprises about a hundred proteins in yeast (Will & Luhrmann, 2011). The spliceosome must correctly recognize the 5’ splice site (5’ SS), the branch-point adenosine (brA), and the 3’ splice site (3’ SS) in order to remove introns (Figure 1a). Correct spliceosome assembly on introns and accurate splice site selection are important, as using the wrong splice sites could ultimately lead to errors in protein translation; erroneous splicing has been linked to cancer in humans and therapeutics designed to modulate splicing are on the horizon (Lee & Abdel-Wahab, 2016). Eight ATP-dependent helicases - comprising three DEAD-box helicases (Sub2, Prp5, Prp28), four DEAH-box helicases (Prp2, Prp16, Prp22, Prp43) and a Ski2-like helicase (Brr2) - ensure splicing fidelity by actively promoting correct spliceosome assembly and splice site usage (Semlow & Staley, 2012). All eight helicases are essential in yeast and conserved in humans (Cordin & Beggs, 2013; Giaever et al., 2002).

**Figure 1:**
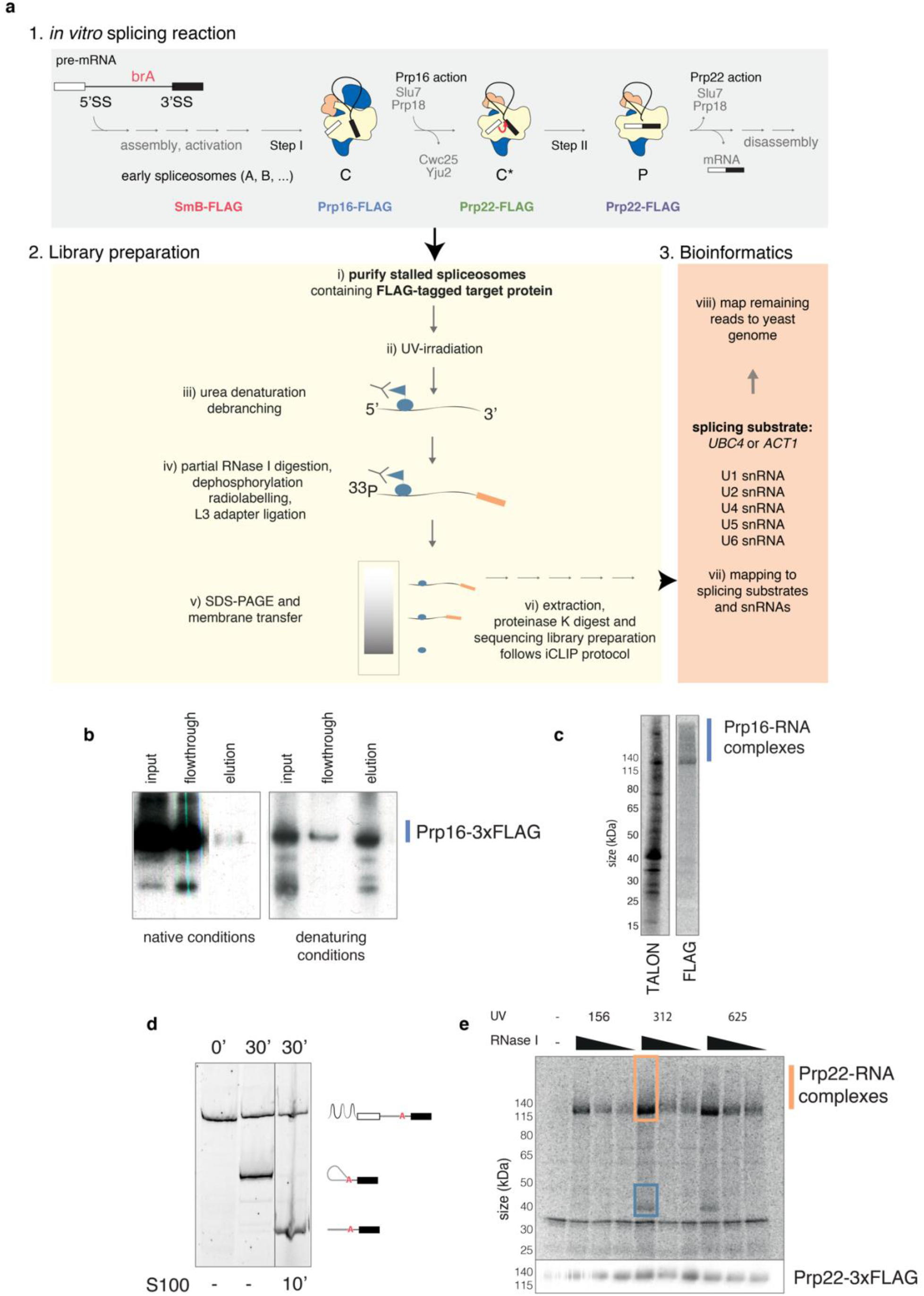
The psiCLIP method reveals the complete RNA binding profile of an RBP in a specific spliceosomal complex. **a)** Spliceosomes are assembled from yeast cell extract on an *in vitro* transcribed pre-mRNA and stalled through different strategies that block the transition towards later stages. The spliceosomes are depicted in schematics, whereas U2 snRNP is coloured in yellow, U5 snRNP in blue, and U6 snRNP in orange. The RNA binding protein of interest is fused to a C-terminal FLAG-tag. After purification of the stalled spliceosome population, spliceosomes are irradiated with a defined UV dose at 254 nm and denatured to release the target protein from the complex and allow FLAG capture of the RNA-protein complex. For spliceosomes stalled at steps containing lariat structures, an additional debranching step was introduced. Controlled RNase I digestion fragments the RNA into suitable pieces for Illumina NGS. The 5’-end is radio-labelled, while a DNA adapter is attached to the 3’-end carrying a sequence for primer annealing for reverse transcription. Finally, the crosslinked RNA-protein complexes are resolved on SDS-PAGE and extracted for RNA-library preparation. Full description can be found in the methods section. **b)** anti-FLAG Western blot detecting Prp16-3xFLAG shows that denaturing conditions are required to capture the tagged protein. Purified spliceosomal complex C containing endogenous Prp16-3xFLAG was captured using anti-FLAG beads after either treatment with urea (right) or under native conditions (left). **c)** Radioblots visualizing RNA after the psiCLIP procedure capturing Prp16 with either TALON beads on His-tag (left) or anti-FLAG beads on 3xFLAG tag. Signal that arises from Prp16 crosslinked to RNA is expected to be larger than the size of Prp16 at 122 kDa. **d)** Treatment of lariats with S100 cell lysate fraction leads to debranching. Intron-lariat intermediates were generated by *in vitro* splicing reactions of 3xMS2-*UBC4*-AC (lane 2). Treatment with S100 for 10 min in debranching buffer leads to full debranching. Only splicing products containing the second exon are visible as the 3’ end of the transcript is fluorescently labelled. **e)** Radioblot for parallel fine-tuning of UV-irradiation and RNase I digest (top panel). Purified complex C* containing endogenous Prp22-3xFLAG was irradiated with the indicated doses of UV (in 100 × μJ/cm_2_), and fragmented by the addition of 5, 0.5 or 0.05 units of RNase I following the psiCLIP procedure as described in the methods section. Bottom panel shows subsequent Western probing with anti-FLAG antibody to visualize Prp22 protein levels. The orange box indicates the optimal condition for Prp22 crosslinking and the membrane area selected for cDNA library preparation. The blue box shows a co-crosslinked protein that was released with the given RNase I concentration.

The DEAD-box helicases Sub2, Prp5, and Prp28 are involved in early assembly of the spliceosome at introns, promoting correct splice site recognition (Cordin & Beggs, 2013). The Ski2-like helicase Brr2 is involved in spliceosome activation prior to the first catalytic step of splicing. Finally, the four DEAH-box helicases Prp2, Prp16, Prp22, and Prp43 remodel the spliceosome through the first (branching) and second (exon ligation) catalytic steps, and coordinate spliceosome disassembly once the reaction has completed (Cordin & Beggs, 2013). Prp16 repositions the pre-mRNA after the branching reaction by removing the branch-helix, formed between the branch point sequence and U2 snRNA, from the active site of the C complex spliceosome (Galej et al., 2016; Semlow et al., 2016; Fica et al., 2017). *In vitro* experiments and structural studies of spliceosomal DEAH-box helicases show a preferred translocation direction from 3’ to 5’ for RNA duplex unwinding (He et al., 2017; Tanaka & Schwer, 2005). Prp16 activity releases the branching factors, which stabilise the branch helix in the C complex, from the spliceosome and generates binding sites for the exon ligation factors Slu7 and Prp18. Prp16 is then replaced by Prp22, which leads to the catalytically active complex C* (Ohrt et al., 2013) and promotes the release of mature mRNA after exon ligation (Company et al., 1991).

Prp22 and the second step factors Slu7 and Prp18 promote exon ligation for most pre-mRNA substrates, during which Prp22 acts in an ATP-independent manner (B. Schwer & Gross, 1998). Exon ligation results in the spliceosomal complex P that still contains both the excised intron and the mature mRNA, which is held by spliceosomal proteins and the U5 snRNA. The mature mRNA is then released from the spliceosome through Prp22’s ATP-dependent activity, in parallel to the release of exon ligation factors Prp18 and Slu7 (Company et al., 1991; Schwer & Gross, 1998; Wagner et al., 1998). Prp22 was shown to unwind duplex RNA *in vitro* in an NTP dependent manner (He et al., 2017; Tanaka & Schwer, 2005). After exon ligation, the helicase was found to interact with the second exon (Beate Schwer, 2008). Therefore it is likely that Prp22 releases the mRNA by translocating along the second exon in a 3’ to 5’ direction, unwinding the U5 snRNA-mRNA duplex (Beate Schwer, 2008).

Despite advances in the structural and compositional analysis of the spliceosome, much is still unknown about the helicase-mediated transitions between spliceosome intermediates (Plaschka et al., 2019). Although previous research indicates the stages at which helicases are required, it is not known exactly where on the snRNA and/or pre-mRNA they bind, how they promote conformational changes in the spliceosome, nor the mechanism by which they ensure splicing fidelity. Both steps of splicing are reversible, offering the potential to prevent suboptimal pre-mRNA transcripts from completing splicing (Tseng & Cheng, 2008). Additionally, DEAD/H-box helicases were proposed to promote the backwards reaction in case a suboptimal site was recognised (Koodathingal et al., 2010). Two proofreading mechanisms are suggested to increase splicing fidelity. (1) The kinetic mechanism posits that the ATPase activity of DEAH-box helicases acts as a timer. Hydrolysis promotes either the productive pathway towards exon-ligation or the discard pathway for suboptimal mRNA transcripts (Semlow & Staley, 2012). The helicase Prp43 for instance was suggested to disassemble spliceosomes in this discard pathway, as the absence of Prp43 helicase led to the accumulation of splicing errors (Mayas et al., 2010). (2) The thermodynamic model proposes that splicing fidelity is enhanced through the stability of competing spliceosome conformations (Semlow & Staley, 2012). Indeed, DEAH-box helicases were proposed to shift the equilibrium between different catalytic conformations of the spliceosome (Query & Konarska, 2004).

Most helicases only transiently bind to the spliceosome, making them difficult to study. Cryo - electron microscopy (cryoEM) structures have shown helicases to be at the spliceosome periphery, but they are detected at much lower local resolutions than other components due to their flexibility. Despite their peripheral location in these structures, helicases can promote remodelling of the RNA-based catalytic core of the spliceosome, but it remains unclear how exactly this occurs.

Additional insights have been gained by site-specific crosslinking of helicases to pre-mRNA substrates in splicing extracts upon incorporation of single photo-reactive residues into the RNA (McPheeters & Muhlenkamp, 2003; McPheeters et al., 2000; Beate Schwer, 2008). However, the identified binding positions were limited to the nucleotides chosen for site-specific modification and were not linked to a specific splicing stage. Single molecule fluorescence resonance energy transfer (smFRET) studies have advanced our understanding of the mechanism of DEAH-box helicases, suggesting they remodel the spliceosome active site from a distance (Semlow et al., 2016). However, this approach does not provide precise positional information about RNA-protein contacts and is technically cumbersome as the positioning of FRET probes must be carefully calibrated for each substrate. Perhaps for this reason, most studies focused on one specific RNA transcript, making it difficult to distinguish transcript-specific effects from general principles.

Transcriptome-wide studies, such as spliceosome iCLIP using antibodies against the SmB proteins (Briese et al., 2019), spliceosome profiling using Prp19-TAP affinity purification (Burke et al., 2018), and Cdc5-GFP + Lea1-Protein A + Snu114-CBP affinity purification (Chen et al., 2018), have uncovered global rules of splicing. However, the contribution of individual splicing factors is difficult to discern in such profiles. In addition, iCLIP of individual splicing factors has revealed their position-dependent capacity to control alternative splicing decisions in mammals (Ule & Blencowe, 2019) but has not led to direct insights into step-specific mechanisms of the core spliceosome. A key issue with these methods is that spliceosomes are not stalled at a specific step, therefore the resulting profiles represent an ensemble of spliceosomal snapshots across many splicing steps, with a bias towards those that are rate-limiting. This prevents us from assigning specific binding sites to individual steps. To understand more detailed rules governing splicing, a method is needed that is capable of identifying RNA contacts formed at defined steps of splicing. Such a method would be particularly useful in resolving the principles that guide the binding of spliceosomal helicases to RNA substrates, in identifying potential transcript-specific effects, and for understanding how the ATPase activities of helicases are linked to their RNA-binding behaviour and thereby to their proofreading activities.

Here, we present psiCLIP (purified spliceosome iCLIP), a method to determine the complete range of contacts between specific splicing substrates and helicases in native spliceosomal complexes of defined states. UV-crosslinking of the enriched complex makes it possible to capture the weak and transient binding of DEAH-box helicases. We used a system of *in vitro* assembled yeast spliceosomes, which allows targeted isolation of helicases at defined points in the splicing pathway and enables functional characterization of helicases using specific mutants. Using these approaches we analysed two DEAH-box helicases, Prp16 and Prp22, which act before and after the second catalytic step respectively (complexes C, C* and P), in the context of multiple RNA transcripts. The high sensitivity and resolution of psiCLIP data provide insights into helicase dynamics and lay the foundation for future mechanistic studies to dissect helicase functions in the spliceosome.

## Results

### psiCLIP enables study of RNA binding proteins in a native spliceosomal complex of a defined state

In addition to conventional iCLIP, which uses crosslinked cells or tissues as starting material (Lee & Ule, 2018), the method has also been adapted to study the binding of purified U2AF2 to pre-mRNAs (Sutandy et al., 2018). Here with psiCLIP, the starting material is yeast cell extract, supplemented with a defined pre-mRNA substrate and for some of our assays, recombinant helicases too. Step-specific spliceosomes are generated by *in vitro* assembly and then purified in a similar way to those used for structural analysis (Wilkinson et al., 2018) (Figure 1a). Spliceosomes are generated by *in vitro* assembly using a specific pre-mRNA substrate (Figure S1), leading to high coverage of sequence reads on the transcript. As in conventional iCLIP, spliceosomes are irradiated with 254 nm ultraviolet light to crosslink proteins with the pre-mRNA substrate and snRNAs. Next, the RNA-binding protein (RBP) of interest is purified under stringent, denaturing conditions, along with any crosslinked RNA, followed by cDNA library preparation and high-throughput sequencing. The computationally processed sequencing reads provide a profile of RBP-RNA contacts that take place within the spliceosome in a specific state.

We used psiCLIP to investigate the binding profiles of the DEAH-box helicases Prp16 and Prp22 that act in the spliceosomal complexes C, C* and P (Figure 1a). After crosslinking, we treated the spliceosome with denaturing conditions, captured 3xFLAG-tagged RBP with anti-FLAG magnetic beads, and eluted under stringent conditions for SDS-PAGE analysis (Figure 1b). Protein-RNA complexes were visualised by radiolabeling the crosslinked RNA, which showed a continuous signal starting around the size of the apo protein towards higher molecular weight, which is expected given the random fragmentation of RNA. Comparison with a His-tag purification via TALON beads demonstrated that the 3xFLAG purification was more specific and therefore we proceeded with 3xFLAG for all experiments in this study (Figure 1c).

To optimise the method, we introduced a debranching step for spliceosomes bound to the lariat RNA structure in order to reduce cDNA truncations at branch-points that could be erroneously interpreted as protein-RNA crosslinks (Figure 1d, Figure S2; Briese et al., 2019; Chen et al., 2018). UV doses were titrated to be efficient but not too high, thus minimising the chance of multiple proximal protein-RNA crosslinks that could bias reverse transcription truncations to the 3’ end of the binding sites (Figure 1e). We also optimised the incubation of purified spliceosomes with a limited amount of RNase to generate RNA fragments with median size of approximately 100nt that are optimal for minimising the sequence preferences of RNase and UV-crosslinking (Haberman et al., 2017; Huppertz et al., 2014). This is particularly important for psiCLIP to ensure that enough sequence is available at the 3’ end of the RNA to allow mapping of the reads. We titrated UV and RNase I doses in parallel (Figure 1e), which showed that a high RNase concentration was optimal for liberating a multi-protein crosslink (blue box) for Prp22 psiCLIP, and that the signal was saturated at 312 × 100 μJ/cm_2_ UV dose. Subsequent steps for cDNA library preparation followed the iCLIP protocol as previously described (Blazquez et al., 2018). The sequencing reads were then analyzed using a reproducible computational workflow, which maps the reads for individual experiments to a custom genomic index containing the pre-mRNA substrate sequences used for the *in vitro* splicing reaction and all spliceosomal snRNAs. Reads that do not map to this custom index are then mapped to the yeast genome and are used for normalisation.

### psiCLIP correctly locates the binding of SmB on snRNAs

To test our protocol, we first chose to examine binding of an integral spliceosomal protein, small nuclear ribonucleoprotein-associated protein B (SmB). The U1, U2, U4 and U5 snRNPs all contain an Sm ring, a heptameric ring structure composed of Sm proteins (E, F, G, D1, D2, D3 and B) around the snRNAs. The Sm binding sites of this ring have been characterized by genetic studies and the U-rich consensus sequence Pu-U(4-6)G-Pu was found to be essential for binding (Guthrie & Patterson, 1988). The Sm proteins have also been modelled in cryoEM structures (Price et al., 1998; Leung et al., 2011; Kondo et al., 2015). However, the exact nucleotide binding positions of the Sm-proteins, including their unstructured tail regions, are unknown for most snRNPs. In order to analyze the SmB binding profile using psiCLIP, spliceosomes were assembled on a substrate with a 3’SS mutation that prohibits progression to the second step of splicing, yielding early splicing complexes and spliceosomes of the first catalytic step (complex C).

The resulting psiCLIP data shows that SmB crosslinks predominantly to snRNAs, with little signal on the pre-mRNA transcript (short *UBC4* shown in Figure 2b-f). We observe crosslinks on all Sm sites, with very few on the U6 snRNA, which is bound by the LSm ring that lacks the SmB protein (Figure 2b-f). There is little signal in the negative controls (no UV/tagged protein; Figure 2b-f, bar graphs). Upon examining individual snRNAs, there is strong psiCLIP crosslinking located within and adjacent to the Sm sites, known to bind SmB (Figure 2g). The three-dimensional structure for the pre-catalytic spliceosomal complex pre-B (Bai et al., 2018) shows that the SmB protein is in close spatial proximity to the U1 snRNA nucleotides detected using psiCLIP (Figure 2h). The 5’ end of U1 snRNA extends into the direction of the SmB protein providing an explanation as to why the crosslinks are mainly found at the beginning of the Sm site motif. Additionally, the crosslinking upstream of the Sm site identified by psiCLIP (G535, A536) are shown in the structure to interact with the long tail of the SmB-protein (Figure 2h). This part of SmB could so far only be modelled into the U1 snRNP in the cryoEM structure of the yeast pre-B complex, due to its high flexibility (Bai et al., 2018). The psiCLIP data suggest that a similar configuration of RNA and SmB protein will be present in the other snRNPs (Figure 2g). This example suggested that psiCLIP could be particularly useful for interactions of flexible protein regions that are challenging for structural methods, and we therefore set out to study the RNA contacts made by transiently acting helicases.

**Figure 2:**
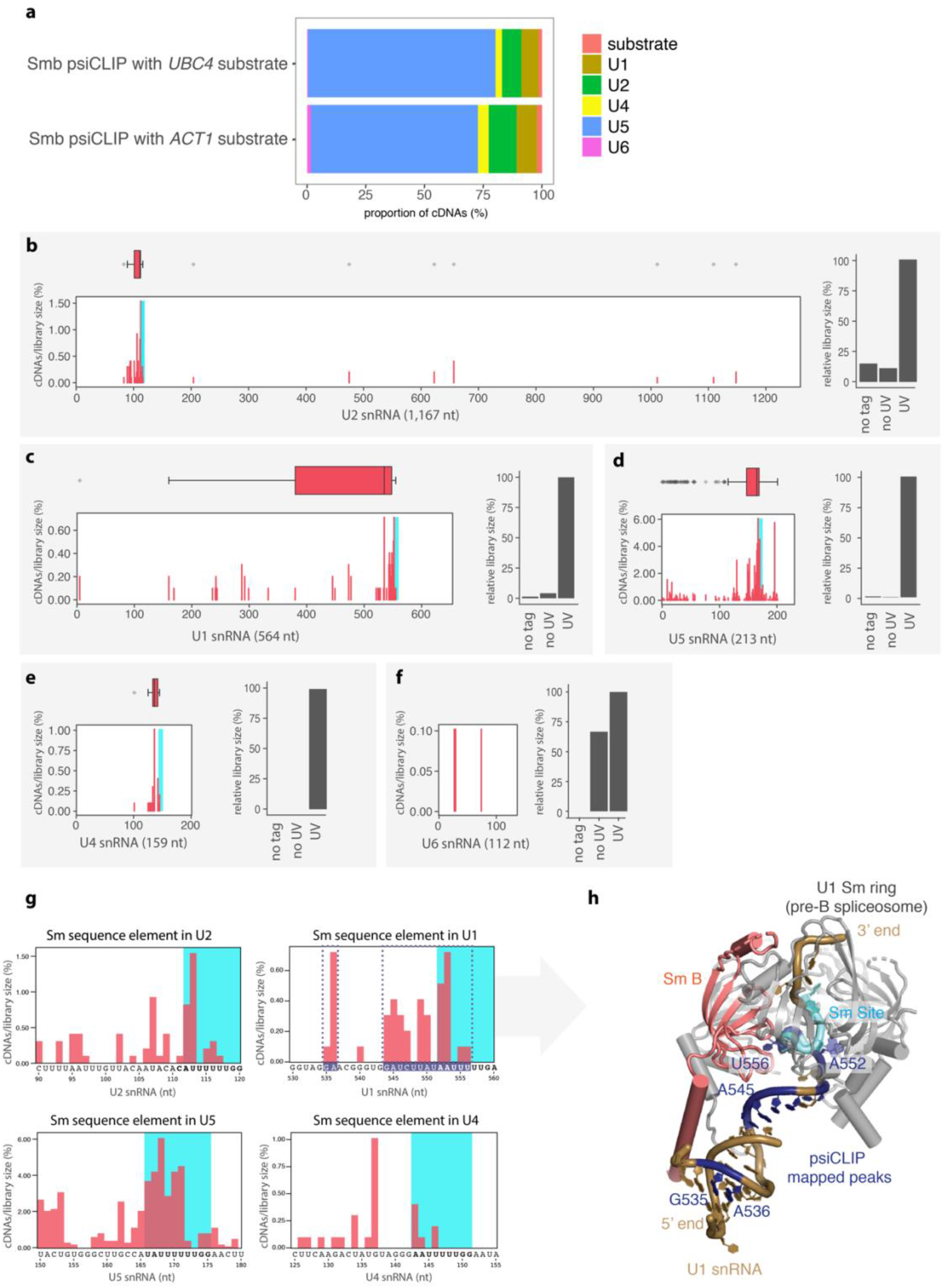
psiCLIP data on SmB mapped onto the snRNAs. **a)** Proportion of cDNAs mapping to substrate and snRNAs for two Smb psiCLIP experiments. **b-f)** Crosslink events (−1 position of cDNA start) are represented as histograms and boxplots aligned to the indicated snRNA. On the histograms, the y-axis indicates the proportion of cDNAs out of all cDNAs mapped to snRNAs and pre-mRNA substrate. On the boxplots, the salmon area denotes the region of highest crosslink density. Due to very low crosslinking on U6 snRNA the boxplot is not shown. The Sm sites are highlighted in blue. The bar charts represent the proportion of cDNAs in samples, normalised to the UV condition, which is shown as 100%. No tag: UV irradiated untagged SmB, No UV: SmB-3xFLAG without UV irradiation and UV: UV irradiated SmB-3xFLAG. **g)** Zoom in to show crosslinking specifically at the Sm sequence elements, which are highlighted in bold letters and blue background. The U2 snRNA region highlighted in purple is shown in G in the crystal structure. **h)** Region containing the U1 snRNP Sm ring of the spliceosomal complex pre-B (Bai et al., 2018) with the SmB protein in salmon, the nucleotides of U1 snRNA in gold, with those showing the highest psiCLIP signal in purple, corresponding to the purple boxes in g. Blue denotes the Sm site, as in g.

### PsiCLIP reveals that Prp16 binding depends on the brA to 3’SS distance

The DEAH-box helicase Prp16 is essential for the transition from complex C, the complex after branching, to the catalytically active complex C* (Ohrt et al., 2013). The helicase binds transiently to the spliceosome to reposition the splicing substrate, removing the brA from the active site, creating space that will later allow the 3’ SS to dock at the same location (B. Schwer & Guthrie, 1992, Wilkinson et al., 2017). This Prp16-dependent spliceosome remodeling causes branching factors Cwc25 and Yju2 to be released and binding sites for the exon ligation factors Slu7 and Prp18 to be formed (Fica et al., 2017; Yan et al., 2016). We performed Prp16 psiCLIP using the purified spliceosomal complex C under three different stalling conditions: (a) alteration of the 3’SS from the canonical AG| to AC| (| marks the splice site) (Galej et al., 2016), (b) addition of recombinant, ATPase-deficient Prp16-G378A mutant (Schneider at al., 2002), or (c) both. To assess substrate-specific effects, Prp16 psiCLIP was performed with *UBC4* or *ACT1* pre-mRNAs. These RNA transcripts were chosen because they have been used extensively in previous spliceosome research.

Despite the additional debranching step, we still observed cDNA truncations at the brA which likely result from truncation at partially digested three-way lariat-junctions (Figure S2), indicating that our debranching of the intron-lariat intermediate was incomplete. Crosslinked proteins nearby may obstruct the site for the debranching enzyme to act on. Therefore we removed reads mapping to a 5nt region around the brA (−2 to +2 around brA) to avoid confounding the analysis. Several observations can be made about Prp16-RNA interactions with the remaining cDNA truncations. First, crosslinks are predominantly found on the pre-mRNA substrate, with little signal on snRNAs (Figure 3a). The snRNAs that are not contained in complex C, such as U1 and U4 snRNA, are under-represented compared with U2 and U5 snRNA that are part of complex C (Figure 3b). Second, Prp16 makes similar RNA contacts regardless of the stalling method, implying high reproducibility of the psiCLIP method and that the three stalling methods result in similar spliceosomal complexes (Figure 3c-d). Third, with both splicing substrates, Prp16 mainly binds downstream of the brA. On *ACT1*, Prp16 mainly binds around 30nt downstream, though binding spans the entire region between the brA and second exon (Figure 3c – sum of all replicates, Figure S3 – three replicates shown separately). On *UBC4*, peak binding occurs around 20nt downstream, and again, the profile spans the full length of the intron from the brA, even extending into the second exon (Figure 3d – sum of all replicates, Figure S3 – three replicates shown separately). These findings include positions previously identified using photo-reactive nucleotides (McPheeters & Muhlenkamp, 2003; Semlow et al., 2016) and confirm inferences from the cryoEM structure, in which 18 nucleotides were predicted to span the distance between the brA and the RNA entry site of Prp16. This prediction was drawn from the six nucleotides closest to the brA which could be modelled into ordered density, with the additional 12 added based on the approximate length that would be needed to reach the RNA entry site of Prp16 (Figure 3e; Galej et al., 2016). Taken together, psiCLIP shows extensive dynamic binding of Prp16 in complex C and provides precise positional information about its binding site for the first time.

**Figure 3:**
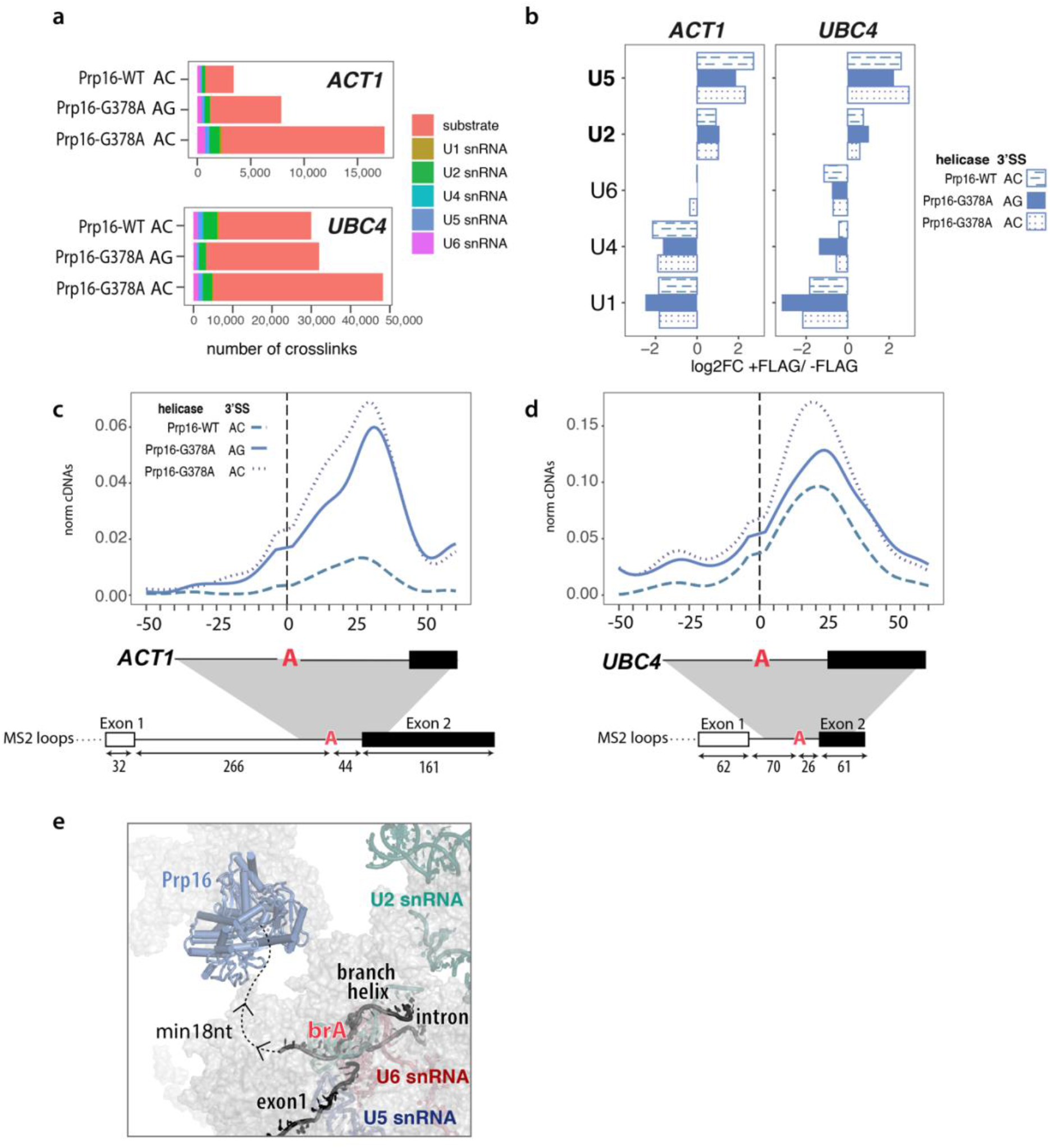
Prp16 psiCLIP data show substrate specific binding in a region downstream of the brA. **a)** Total number of crosslinks summed across three replicates for each condition (UV crosslinked, 3xFLAG-tagged samples). Colour denotes the RNA species the crosslinks map to. **b)** log2 fold change enrichment of snRNAs in FLAG-tagged Prp16 vs. untagged samples. Note that for this analysis, crosslinks were first normalised to the total number of snRNA crosslinks in each sample. **c, d)** Mapping of Prp16 psiCLIP data onto *ACT1* and *UBC4* splicing substrates. Three replicates were summed. The smoothed lines show the truncation events normalized to library size, with the control signal subtracted from the UV signal. Lines were Gaussian smoothed with a window size of 20nt. Positions along the transcript are shown relative to the brA. Crosslinking signal derived from the three different stalling methods (substrate with 3’SS mutation / dominant negative Prp16 / combination of both) result in the same pattern. The main peak that reflects the position of the Prp16 helicase is slightly further downstream on *ACT1* than on *UBC4*. **e)** CryoEM structure of Prp16 in C complex shows the helicase at a distance from the brA. Six nucleotides were built into the density after the brA (brA +6 is the last ordered nucleotide). The dotted line gives an indication of where the intron and the second exon of the transcript could expand to. A minimum distance of 18nt between the brA and the entry site of Prp16 was estimated in (Galej et al., 2016).

To test the constraints on Prp16 binding, we designed transcripts based on *ACT1* with increasing lengths between the brA and 3’ SS. Randomised 20 and 40 nucleotide sequences with the same GC content as the *ACT1* intron were inserted after the brA and were shown to splice with comparable efficiency to the original transcript (Figure 4a-d). psiCLIP was performed with the spliceosomal complex C assembled onto these substrates and stalled using the ATPase-deficient Prp16 mutant (Prp16-G378A). As the distance between brA and 3’SS was increased, the binding region of Prp16 also expanded to fill this new space (Figure 4e-h). We observed crosslinking about 30nt upstream of the brA, but this was disregarded as it was inconsistent between replicates and it was also detected in the control samples (Figure S4). Taken together, these results indicate that Prp16 initially binds downstream of the brA and contacts the substrate wherever it is accessible as a single strand. Most likely, the longer the sequence between the brA and the 3’ SS, the more accessible RNA is available. Since broad contacts are observed with both the mutant, which has diminished translocation activity indicated by a residual ATPase activity (Schneider at al., 2002), and the WT Prp16 helicase (Figure 3 c,d), it remains unclear if the broad binding reflects movement during translocation or other forms of dynamic contacts, such as binding, release and re-binding.

**Figure 4:**
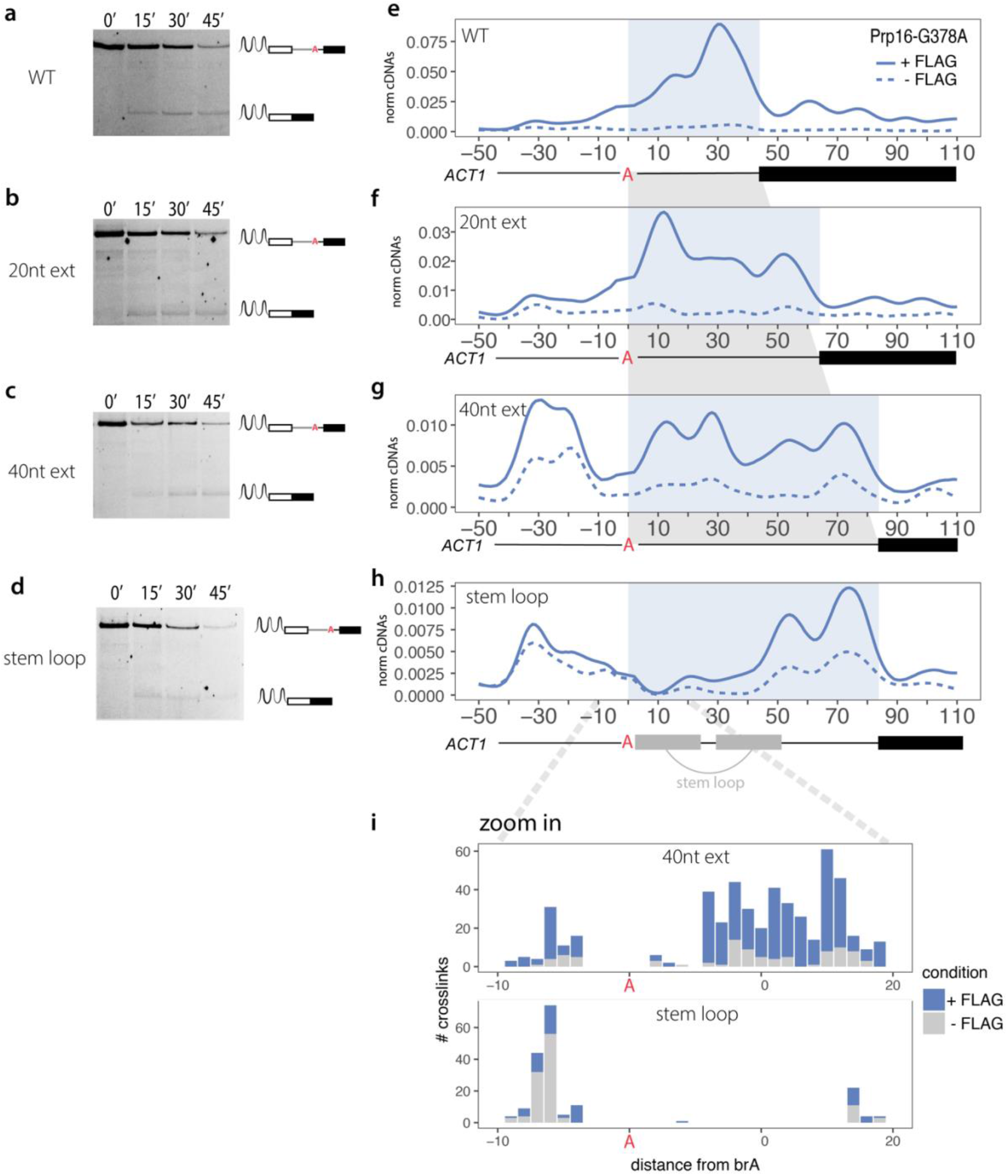
Prp16 psiCLIP binding window expands to fill space between brA and 3’SS in *ACT1* extended substrates. **a-d)** Denaturing RNA PAGE after *in vitro* splicing reactions with fluorescently labeled substrates shows that all versions of the extended *ACT1* substrate are capable of being spliced. Only splicing products containing the second exon are visible as the 3’ end of the transcript is fluorescently labelled. **e-h)** Mapping of the psiCLIP data generated using Prp16-G378A mutant protein and various forms of the *ACT1* splicing substrate including artificial extensions between the brA and the 3’SS. The splicing substrates are aligned onto the brA, and the distance between the brA and the 3’SS is highlighted in light blue. The smoothed lines show the truncation events normalized to library size, with the control signal subtracted from the UV signal. Lines were Gaussian smoothed with a window size of 20nt. Positions along the transcript are shown relative to the brA. The crosslinking signal spreads across the whole length of the sequence between the brA and 3’SS. **i)** zoom in on region around the brA for ext40 and stem loop conditions to show the complete loss of crosslinking downstream in the stem loop experiment.

The branch-helix forms between U2 snRNA and the region upstream of brA on the substrate (Figure 3e). A long standing puzzle has been how Prp16, transiently located at the spliceosomal periphery, can remodel this branch-helix that is buried within the spliceosome. In order to make exon ligation happen, the branch helix has to be moved out of the catalytic core to make space for the second exon to come in. To address this question, we introduced a 40nt stem-loop into *ACT1*, with the intention of creating an obstacle between Prp16 and the brA (Figure 4d,h). This substrate was capable of being spliced, indicating that Prp16 action is not meaningfully hindered by the inserted sequence (Figure 4d). Performing psiCLIP, we observed a 3’ shift in Prp16 crosslinking (Figure 4e), such that the 5’ arm of the stem loop showed hardly any crosslinking signal and the 3’ arm a reduced crosslinking signal; this suggested that the sequence indeed formed a secondary structure that hindered Prp16-binding. Crosslinking was seen towards the 3’ end of the 3’ arm (Figure 3E); since RBPs preferentially crosslink to single-strand RNA (Rogelj et al., 2012) this suggests that either the stem loop was not completely formed and allowed Prp16-binding in that region or that the helicase mutant unwound it partially with its residual ATPase activity. On the assumption that most of the stem loop stays intact during the remodelling event, this result is consistent with a model in which Prp16 acts on the branch helix without translocating through the brA in order for splicing to occur (Semlow et al., 2016). That said, it is plausible that the WT helicase would be able to unwind and translocate through the stem loop, however using the mutant was required here to achieve the stalling necessary to catch the helicase.

Taken together, our findings demonstrate that psiCLIP has a large dynamic range to test hypotheses about helicase remodelling in the context of intact native spliceosomes, thus providing complementary functional assays to FRET and cryoEM studies.

### Prp22 psiCLIP detects repositioning of the helicase after the second step of splicing

After Prp16 is released, leading to the formation of the spliceosomal complex C*, which is competent for exon ligation, Prp22 binds the spliceosome at a similar location on Prp8 (Figures 1, 3e and 5a; Fica et al., 2017; Galej et al., 2016). Exon ligation results in the spliceosomal complex P that still contains both the excised intron and the mature mRNA. RNA density observed in the Prp22 helicase core in the cryoEM structure of complex C* was assigned to the second mRNA exon, with approximately 16 nucleotides needed to span the distance between the last nucleotide of the first exon and the center of the Prp22 RNA binding pocket (Fica et al., 2017). However, biochemical experiments suggested Prp22 binds on the intron close to the 3’ SS before exon ligation (McPheeters et al., 2000). In the stalled spliceosomal complex P, structural analysis found Prp22 positioned between nucleotide 14 to 21 on the second exon of the *UBC4* transcript (Wilkinson et al., 2017). Thus the precise position of Prp22 on the substrate before and after exon ligation remained unclear.

**Figure 5:**
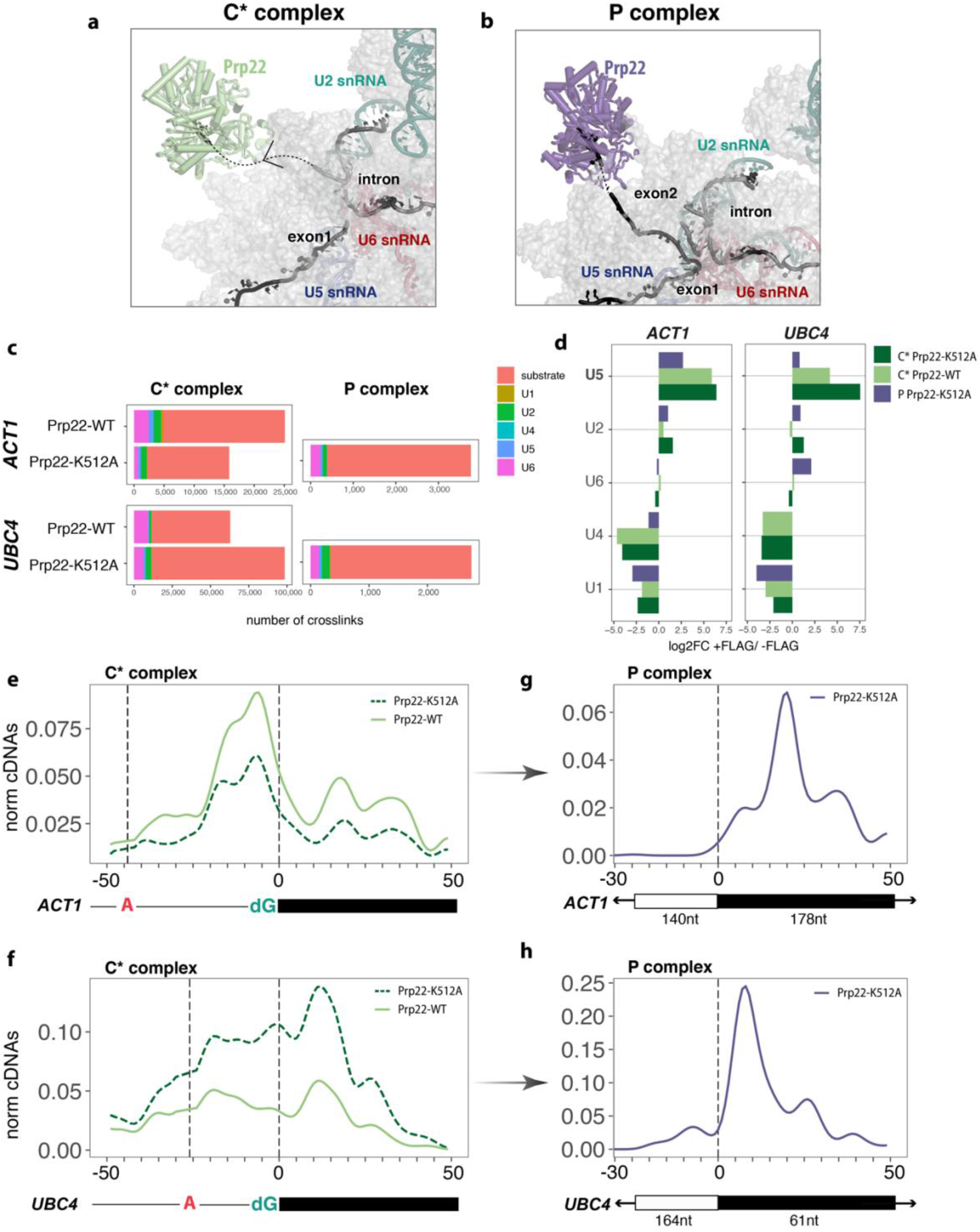
Prp22 psiCLIP indicates an extensive binding profile in C* complex, and this shifts to a narrow peak of binding in P complex. **a,b)** Cryo-EM structure of Prp22 in C* complex and P complex shows the helicase in green and purple, respectively. The dotted line marks regions of RNA that were not built and are just predictions. **c)**Total number of crosslinks summed across two UV crosslinked, FLAG-tagged replicates for each condition. Colour denotes the RNA species the crosslinks map to. **d)** log2 fold change enrichment of snRNAs in FLAG-tagged vs. untagged samples. Note that for this analysis crosslinks were first normalised to the total number of snRNA crosslinks in each sample. **e-h)** Mapping of Prp22 (WT and mutant) psiCLIP data onto *ACT1* and *UBC4* transcripts. The smoothed lines show the truncation events normalized to library size, with the control signal subtracted from the UV signal. Crosslinks are shown after Gaussian smoothing with a 10 nt window. Positions on the substrate are given relative to the 3’SS. P complex crosslinks are shown with the intron mapping reads removed. **e)** Prp22-WT and Prp22-K512A psiCLIP on *ACT1* substrate in C* complex. **f)** Prp22-WT and Prp22-K512A psiCLIP on *UBC4* substrate in C* complex. **g)** Prp22-K512A psiCLIP on *ACT1* substrate in P complex. **h)** Prp22-K512A psiCLIP on *UBC4* substrate in P complex.

We performed Prp22 psiCLIP using multiple splicing substrates after enriching for spliceosomes stalled either before (complex C*) or after exon ligation (complex P). We enriched for complex C* by pausing splicing with *UBC4* and *ACT1* transcripts in which the 3’SS G is replaced with deoxynucleotide dG, preventing the second transesterification reaction. psiCLIP was then performed using both the wild-type and the dominant negative Prp22-K512A ATPase-deficient mutant. Again, there was little signal on snRNAs, with under-representation of U1 and U4 snRNAs which are absent in C* complex (Figure 5c,d). Prp22 binding covers a broad region around the 3’ SS on both *UBC4* and *ACT1* transcripts. The region starts just around the brA, and extends far into the second exon, meaning that this part of the transcripts is accessible for the helicase to bind. It includes the 3’SS −8 crosslink position that was previously identified using a second step-deficient substrate in splicing extract (McPheeters et al., 2000). Both the wild-type and mutant Prp22 show a similar psiCLIP profile, confirming that binding is ATP-independent, as suggested previously from structural work, where RNA density next to an open conformation of the helicase is found without ATP (Figure 5e,f; Fica et al., 2017).

Next, we performed Prp22 psiCLIP after enriching for spliceosomal complex P by adding recombinant Prp22-K512A dominant negative mutant protein, which prevents the release of ligated exons. P complex psiCLIPs were enriched for junction reads, with 10-15% of mapped reads containing the canonical splice junction, compared with a maximum of 0.0003% for the C* complex (Figure S5). We found sharp binding peaks in the second exons of both splicing substrates at +10 and +20nt downstream of the exon-exon junction in *UBC4* and *ACT1* respectively, with binding extending as far as +35nt (Figure 5g,h). These results agree with the structural analysis of complex P in which Prp22 was suggested to bind between nucleotides 14 and 21 on the second exon of the *UBC4* transcript (Wilkinson et al., 2017). We also identify positions +10, +17, +22 and +35nt on the *ACT1* second exon reported by site-specific crosslinking (Beate Schwer, 2008). Thus Prp22 binds at similar positions within the second exon both before and after exon ligation, but binding to the intron and the start of the second exon are dramatically decreased after exon ligation in the P complex (Figure 5g,h).

### PsiCLIP of Prp22 is sensitive to the presence of auxiliary factors

Exon ligation of *ACT1* and *UBC4* requires the second-step factors Slu7 and Prp18 (James et al., 2002). It has been suggested that Slu7 binds the spliceosome first, followed by Prp18 and then Prp22 (James et al., 2002). In complex P, Prp18 binds near the 3’SS and stabilizes its docking in the active site (Wilkinson et al., 2018). Prp18 was also suggested to promote Prp22 binding to the spliceosome (James et al., 2002), yet the structures of complexes C* and P did not identify direct contacts between Prp18 and Prp22. We therefore sought to investigate whether Prp18 affects Prp22 binding patterns by performing psiCLIP upon depletion of Prp18 from splicing extracts. We also compared the WT and ATPase-deficient Prp22-K512A to assess how Prp22 binding pattern depends on its ATPase activity.

As expected, the depleted extract accumulates lariat intermediate regardless of Prp22 status, indicating a defect in the second catalytic step; this was rescued by adding recombinant Prp18 (Figure S6a; James et al., 2002). Next, psiCLIP was performed using *ACT1* transcripts containing either the canonical or dG modified 3’SS. Purified complexes contained lariat-intermediate and thus appeared stalled after the first catalytic step (Fig. S6b,c). We observed a striking difference in the transcript binding of WT and mutant Prp22 helicases in Prp18- deficient spliceosomes. The WT Prp22 psiCLIP displayed very little binding in the Prp18- deficient spliceosomes (Figure 6b,c, Figure S6b-f), whereas the ATPase-deficient Prp22 helicase showed similar binding in the presence (complex C*) or absence of Prp18 (Figure 6a,b,c). In contrast to WT Prp22, the ATPase-deficient mutant Prp22-K512A allows more exon-ligation at the canonical splice site, as indicated by the presence of spliced mRNA reads (Figure S6g). Indeed, some spliceosomes clearly reached the exon-ligation conformation as evidenced by the pronounced Prp22 peak observed in the second exon, which is indicative of P complex Prp22 binding and is enriched on the canonical substrate vs. the dG substrate (asterisk, Figure 6c).

**Figure 6:**
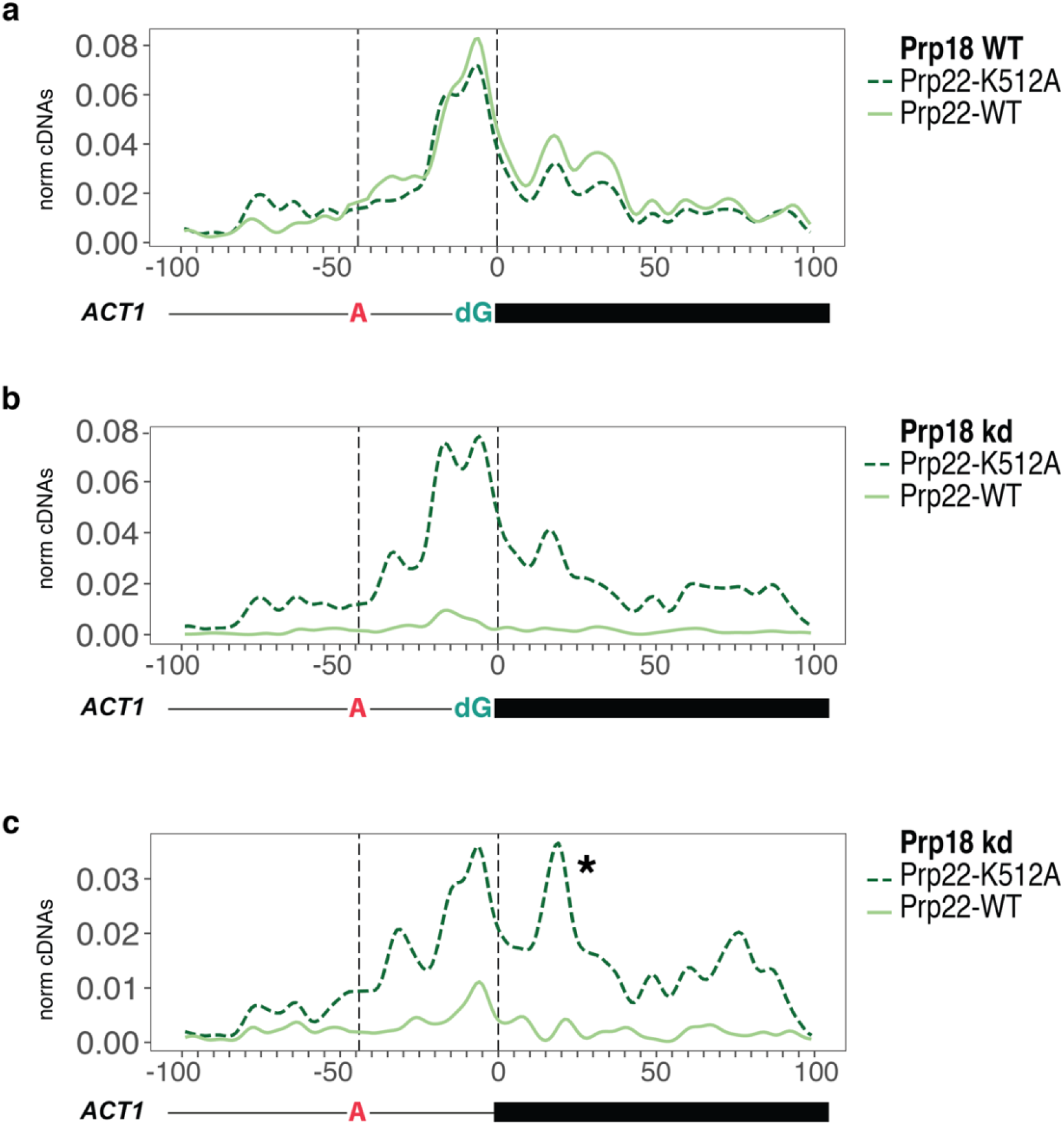
Prp18 depletion abrogates wild-type Prp22 binding on pre-mRNA substrate, but <u>not</u> the binding of ATPase deficient mutant Prp22. Prp22 binding upon Prp18 depletion. Crosslinks are shown as normalized to the sum of all yeast genome mapped cDNAs, with untagged signal subtracted from tagged and a 10nt window gaussian smooth. **a)** Prp22 psiCLIP without Prp18 depletion on an *ACT1* substrate with dG mutation. Both WT and ATPase mutant Prp22 are shown. **b)** Prp22 psiCLIP under Prp18 depletion on an *ACT1* substrate with dG mutation. Both WT and ATPase mutant Prp22 are shown. **c)** Prp22 psiCLIP under Prp18 depletion on an *ACT1* substrate with wild-type AG 3’SS. Both WT and ATPase mutant Prp22 are shown. *represents P complex Prp22 binding signature.

Nonetheless, the majority of the psiCLIP signal derives from lariat-intermediates, suggesting that binding of wild-type Prp22 is destabilized upon Prp18 depletion, in agreement with previous biochemical data (James et al., 2002). Thus psiCLIP is sensitive to disruption in the composition of the spliceosome.

## Discussion

Splicing fidelity is safeguarded by eight ATP-dependent helicases, but the exact positions of their interactions with snRNAs and the pre-mRNA remain poorly understood. This knowledge is required to understand how they promote conformational changes in the spliceosome and ensure splicing fidelity. We developed psiCLIP to gain insights into the detailed binding patterns and molecular functions of two of the most elusive helicases - Prp16 and Prp22 - at defined spliceosomal states. The method is highly reproducible and defines the precise positions of contacts between a helicase and its RNA substrate.

We employed psiCLIP to tackle three key questions about RNA helicase functions. First, we asked how Prp16, located at the periphery of the spliceosome, can reposition the splicing substrate in the catalytic core after the branching reaction. Surprisingly, instead of binding at a fixed distance from the brA, we found that Prp16 binds over the full space between the brA and 3’SS. This indicates that Prp16 initially binds downstream of brA, guided by its interaction with Prp8, and can subsequently translocate and potentially re-bind along all the rest of the intron that is available as a single stand. Elegant single molecule FRET data proposed that Prp16 might act via a winching mechanism, such that its translocation movement is physically stopped by the spliceosome body and thereby transferred into a pulling action that disrupts RNA structures in the active site so that the branch helix is moved out of the active site (Semlow et al. 2016). Our Prp16 psiCLIP data shows that Prp16 binding, and thus presumably translocation, can extend into unstructured RNA inserted after the branch helix. By contrast, Prp16 could not appreciably bind into a structured stem loop region, suggesting that it cannot disrupt strong RNA helices. Despite this, all of our extended constructs spliced. Our data thus provide support for a mechanism of action at a distance in which Prp16 winches towards the branch helix but remodels the spliceosomal active site without disrupting the branch helix.

Second, we asked how Prp22 binding changes during the transition from complex C* to P, and how this depends on Prp22 ATPase activity. The 3’SS was thought to be in the spliceosomal catalytic centre ready for ligation in complex C* (Will & Luhrmann, 2011). However we detected widespread Prp22-binding around the 3’ SS, spreading upstream into the intron, indicating that a large portion of spliceosomes captured in this state don’t exhibit a stably docked 3’ exon, consistent with the absence of a stably docked 3’SS in the cryoEM structure of the C* complex. The extended binding profile in the C* complex suggests that in the absence of a stably docked 3’SS Prp22 can translocate into the intron region of the lariat-intermediate and potentially reject these spliceosomes, thus ensuring that exon ligation occurs only at the correctly docked 3’SS. The strong signal observed for the wild-type Prp22 protein as well as the broad profile seen for the ATPase-deficient K512A mutant suggest multiple rounds of binding and dissociation, rather than a single ATP-dependent translocation and dissociation event. Notably, the psiCLIP binding profile is maintained upon inhibition of Prp22 ATPase activity, consistent with an ATP-independent role for Prp22 in exon ligation (B. Schwer & Gross, 1998). In complex P, Prp22 binding shifts to a narrower region in 3’ exon, and this exonic binding pattern is similar between complex C* and complex P. From this position, Prp22 will start to release the spliced mRNA from the spliceosome. The repositioning of Prp22 observed after exon ligation in the P complex cannot be explained by a processive helicase translocation, as the helicase proceeds in a 3’ to 5’ direction. Instead, we speculate that more than one Prp22 molecule is required per spliceosome for the transition from complex C* to P, or that the helicase releases and rebinds the RNA transcript several times during this transition. Thus, the more narrow binding to the exon in P complex may be coupled to stable docking of the 3’SS and exon-ligation, allowing Prp22 to sense completion of catalysis and release the mRNA in an ATP dependent manner.

Finally, we asked how the exon-ligation factor Prp18 might influence Prp22 binding. We showed that substrate binding by Prp22 is destabilized when Prp18 is absent. As expected, the Prp22 ATPase-deficient mutant remained bound to both the canonical and dG substrate upon Prp18 depletion, and under the canonical condition we detected abundant exon-exon spanning reads corresponding to ligated exons (Figure S6g). This indicates that when Prp22 ATPase activity is compromised splicing can proceed in the absence of Prp18, which would agree with an earlier study in which Prp22 ATPase activity at this stage was only shown to be required for rejecting suboptimal splice sites after branching to prevent mis-spliced exon-ligation (Semlow et al., 2016). Strikingly, binding by the wild-type Prp22 was significantly reduced in the absence of Prp18, consistent with a model in which Prp22 dissociates from these complexes and cannot rebind. This interpretation is supported by structural studies showing that the absence of Prp18 destabilizes the C* and P conformation (Wilkinson et al., 2017) and would likely weaken Prp22 binding onto Prp8, which may be necessary for stable association with the substrate. In contrast, the ATPase-deficient K512A mutant is intrinsically less prone to dissociation, thus explaining the strong signal observed even in the absence of Prp18. Thus, our data would also be consistent with a role for Prp22 ATPase activity in rejecting incompletely assembled spliceosomes. Our psiCLIP data provides valuable information about active and passive helicase activities and lends new mechanistic support for Prp22 proofreading in preventing erroneous exon-ligation.

Further biochemical work will be required to fully elucidate the mechanisms behind the phenomena uncovered by psiCLIP. While psiCLIP is well suited for comparisons between conditions, the absolute crosslinking levels in psiCLIP are affected by the variable crosslinking efficiencies of nucleotides and amino acids that can lead to biases at the sequence level, as in all techniques utilizing UV crosslinking. It is known that uridine is the most favorable crosslinker, followed by guanosine, cytosine and adenosine respectively (Sugimoto et al., 2012). This limitation is important to take into consideration when making comparisons between transcripts and/or proteins. Looking to the future, we envision that this method can be extended to study helicases and other RBPs in additional biochemically defined RNP complexes, such as the RBP-rRNA interactions during ribosome biogenesis. Finally, time-resolved psiCLIP of helicases could be used to track the binding profile of a helicase in motion during RNP remodeling. Thus, it is clear that psiCLIP can help resolve fundamental mechanisms of RNP dynamics and remodeling. With this study, we hope to have made a complementary contribution towards Kiyoshi Nagai’s long commitment to understanding spliceosome function through structure.

## Acknowledgements

We thank Benjamin Porebski for sequencing the cDNA libraries and Sebastian Fica for a generous gift of reagent. We also thank Clemens Plaschka and Flora Lee for critical reading of the manuscript. This work was supported by funding from the Medical Research Council (MC_U105184330), the European Research Council under the European Union’s Seventh Framework Programme (FP7/2007-2013) / ERC grant agreement [617837]. L.M.S. was supported by a Boehringer Ingelheim Fonds Fellowship. This work was supported by the Francis Crick Institute which receives its core funding from Cancer Research UK (FC010110), the UK Medical Research Council (FC010110), and the Wellcome Trust (FC010110). N.M.L is a Winton Group Leader in recognition of the Winton Charitable Foundation’s support towards the establishment of the Francis Crick Institute. N.M.L and J.U are additionally funded by a Wellcome Trust Joint Investigator Award (103760/Z/14/Z), and N.M.L receives funding from the MRC eMedLab Medical Bioinformatics Infrastructure Award (MR/L016311/1) and core funding from the Okinawa Institute of Science & Technology Graduate University.

## Author Contributions

L.M.S. established the method psiCLIP, with advice from M.H. and J.U.. L.M.S. and C.C. designed psiCLIP experiments for SmB, Prp16 and Prp22. L.M.S. generated yeast strains, purified the helicase mutant proteins and performed the psiCLIP experiments. L.M.S. and A.J.N. prepared the splicing substrates and performed the biochemical experiments. A.J.N. cloned, expressed and purified Prp18. S.M.F., A.J.N. and C.O. contributed to the project through their knowledge and experience of yeast splicing. C.C. performed all computational analysis. C.N. supported the generation of yeast strains and optimized debranching steps. The manuscript was written by L.M.S and C.C., with input from all authors. K.N., J.U. and N.L. initiated and coordinated the project.

## Methods

### Yeast strains

All yeast strains used in this study are listed in Table 1. BCY123 (*MATa, can1, ade2, trp1, Ura3-52, his3, leu2-3, 112, pep4::his+, prb1::leu2+,bar1::HisG+, lys2::pGAL1/10-GAL4+*) is the parent strain for all modifications. In brief, sequences coding for protein tags and the respective resistance cassette were amplified with about 60 nt homology to the end of the target gene and the beginning of its 3’ UTR. The linear PCR product was transformed into BCY123 and the cells were plated on selective media containing either 100 μg/mL ClonNAT, 300 μg/mL hygromycin, or 250 μg/mL G418. Positive clones were verified by Sanger sequencing and expression of the tagged protein was analyzed by western blot using antibodies against the tag. Strains containing several C-terminal tags underwent the cycle several times.

**Table 1:** yeast strains included in the study. First column gives the modification compared to the parent strain BCY123. For strains with a genomic integration (indicated column 3), the resistance gene that was used for integration of the C-terminal tag is indicated with its promoter and terminator sequence. Plasmids for overexpression of recombinant proteins are given in column 3 together with the respective selection marker.

| strain | Resistance cassettes | integration |
| --- | --- | --- |
| SmB-3xFLAG-His8 | TEF:nat:TEF | genomic |
| Prp16-3xFLAG-His8 | TEF:nat:TEF | genomic |
| Prp22-3xFLAG-His8 | TEF:nat:TEF | genomic |
| Prp22-3xFLAG,<br>Prp18-3xHA,<br>Slu7-9xcmyc | TEF:nat:ADH1,<br>TEF:kanMX:TEF,<br>TEF:hph:scCYC1 | genomic |
| Prp18-3xHA,<br>Slu7-9xcmyc | TEF:kanMX:TEF,<br>TEF:hph:scCYC1 | genomic |
| Prp16-G378A-CBP-3xFLAG-His6 | URA, TRP | Plasmid: pRS424,<br>pRS426 |
| CBP-Prp22-K512A-3xFLAG-His6 | URA, TRP | Plasmid: pRS424,<br>pRS426 |

For recombinant protein expression, plasmids based on pRS424 and pRS426 vectors carrying *TRP1* and *URA3* markers, respectively were transformed into BCY123 (Table 1). Protein expression is under control of the GAL-GAPDH hybrid promoter. Construct modification by Kunkel mutagenesis (Kunkel, 1985) was verified by Sanger sequencing.

### Preparation of splicing substrates

For in vitro transcriptions, the desired sequence was cloned under the T7 promoter into a pUC-based vector (Table 2) containing a 3xMS2 stem loop sequence either at the 3’ or 5’ end (Zhou et al., 2002). To generate artificial extensions of the *ACT1* substrate, 20 nt long oligonucleotides with a random sequence were generated keeping the GC content the same as in the area between brA and 3’ SS and cloned into a restriction site following the brA position. One clone each with a single 20 nt insertion (5’-CGA TTT TAT TTA TTT GAT CT-3’), a 40 nt tandem insertion (5’-CGA AAA TCA AGA TAA ATA ATC GAA AAT CAA GAT AAA TAA T-3’) and a 40 nt head-to-head insertion for an RNA stem loop generation (5’-CGA TTA TTT ATC TTG ATT TTC GAA AAT CAA GAT AAA TAA T-3’) was selected. Long RNA pieces were generated by run-off transcription.

**Table 2:**
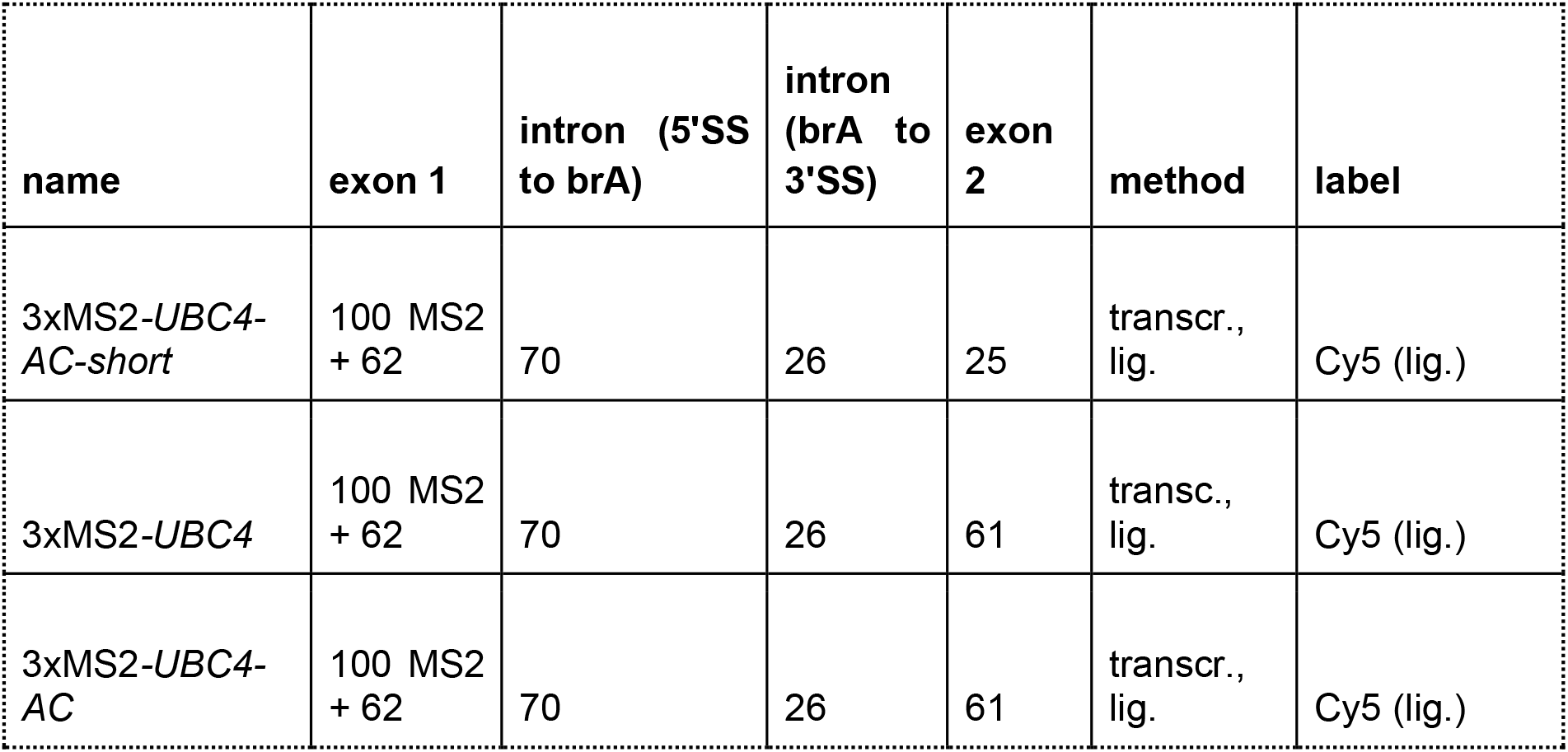
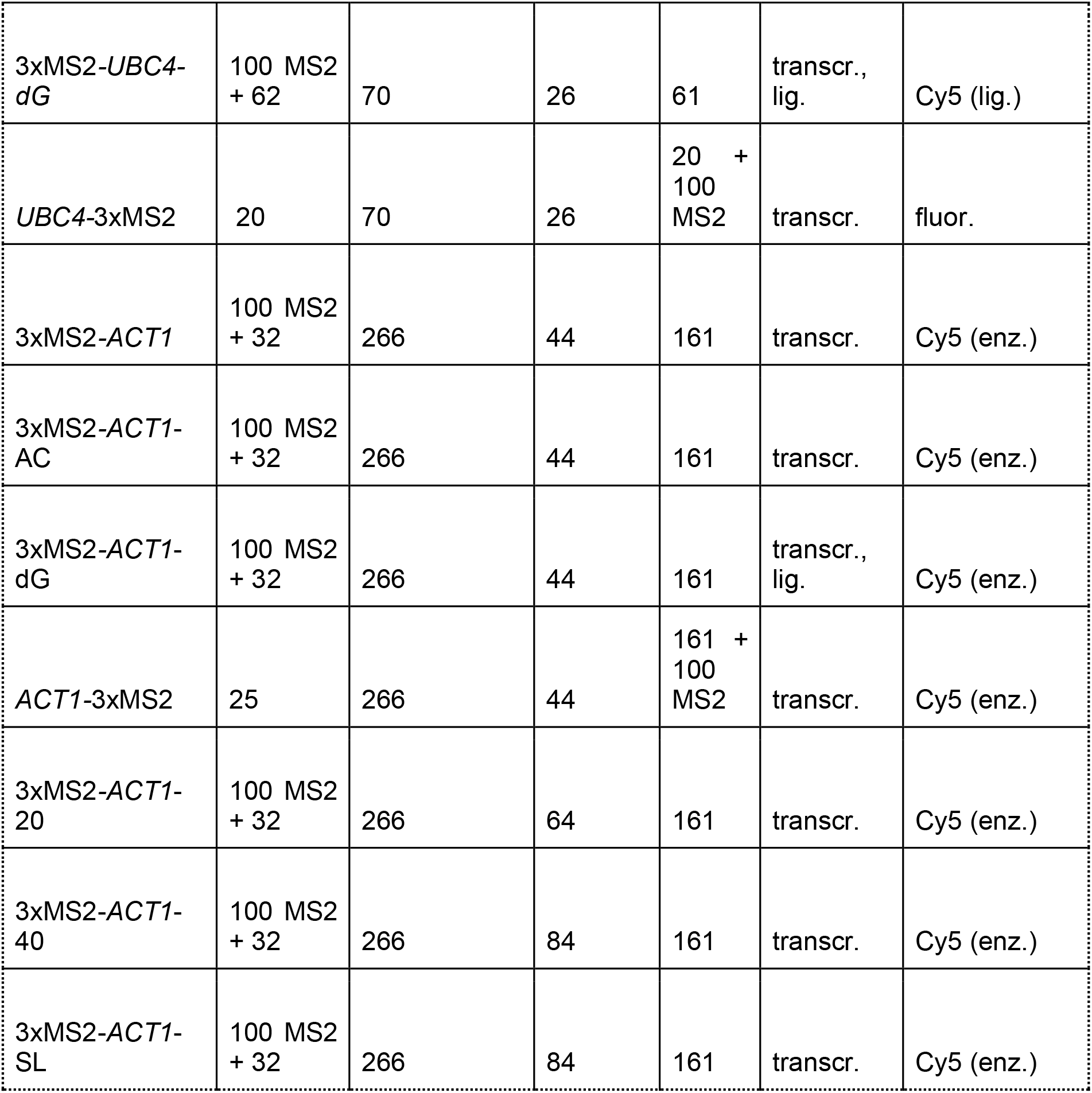
Splicing substrates. First column gives the name including from which gene the transcript derived. The next five columns give the length between the recognition sequences (5’SS / brA / 3’SS) as well as the total length. All lengths are given in nt. Finally, the method via which the transcript was produced (method) and fluorescence-labelled (label) is indicated.

Some splicing substrates, e.g. substrates containing a deoxy-nucleotide, were generated by ligation from a long 5’ piece ending before the 3’ splice site and short 3’ oligonucleotides. For *UBC4* substrates, oligonucleotides carrying a 3’ Cy5 fluorophore (purchased from Sigma) were ligated subsequently. For *ACT1* substrates, both an oligonucleotide and another RNA piece that was generated by run-off transcription were ligated subsequently. To generate precise 3’ or 5’ ends for ligation, these transcripts included a hepatitis delta virus or hammerhead ribozyme sequence at the respective end. RNA pieces were joined by splint-mediated ligation using T4 DNA ligase, essentially as described in (Moore & Sharp, 1992), and ligated transcripts gel purified after each ligation step. Fluorescent labels were attached to splicing transcripts by chemical or enzymatic methods for fluorescein or Cy5-labelling (Table 2).

### Recombinant protein preparation from yeast

BCY123 cells were co-transfected with expression vectors pRS426 and pRS424. Positive transformants were grown in 24 L YM4 selective media supplemented with 1% raffinose at 30°C. Protein expression was induced with 2% galactose (final concentration) at OD600 1.0. After 12-16 hours further growth at 30°C, cells were harvested, resuspended in 1 volume 2 × lysis buffer (2 M NaCl, 100 mM Tris-Cl pH 9.0, 2 mM imidazole, 20 mM β-mercaptoethanol (β-ME), 0.2 % IGEPAL CA-630, 4 mM CaCl2, 2 mM magnesium acetate, EDTA-free protease inhibitor cocktail (Roche)), and frozen in liquid nitrogen in droplet form. Cells were disrupted in a 6870 Freezer/Mill (SPEX SamplePrep) under the following settings: 4 cycles, 2 min precool time, 2 min run time, 1 min cool time between cycles and a rate of 12 cps. After thawing, the pH of the extract was raised to 8.5 by addition of 1M Tris base. Cell debris was removed by ultracentrifugation at 195,000 × g for 90 min. The supernatant was incubated with 2 mL Calmodulin-sepharose beads (home made) for 12-16 hours at 4°C. Beads were washed with 5 × 50 mL CAL wash buffer (500 mM NaCl, 20 mM Tris-Cl pH 8.0, 2 mM CaCl2, 1 mM magnesium acetate, 1 mM imidazole, 10 mM β-ME) and the proteins eluted in 10 × 5 mL CAL elution buffer (500 mM NaCl, 20 mM Tris-Cl pH 8.0, 2 mM EGTA, 1 mM magnesium acetate, 1 mM imidazole, 10 mM β-ME). Protein-containing fractions were pooled and dialyzed against Ni-NTA binding buffer (1 M NaCl, 20 mM Tris-Cl pH 8.5, 5 mM imidazole, 10 mM β-ME) for 4 h at 4 °C. After 14 h binding to 4 mL Ni-NTA agarose beads (Qiagen), the beads were first washed with 15 mL Ni-NTA binding buffer and then with 15 mL Ni-NTA wash buffer (1 M NaCl, 20 mM Tris-Cl pH 8.5, 15 mM imidazole, 10 mM β-ME). The protein was eluted in about six 4 mL fractions with Ni-NTA elution buffer (1 M NaCl, 20 mM Tris-Cl pH 8.5, 250 mM imidazole, 10 mM β-ME). Helicases were dialyzed against the respective helicase storage buffer for 4 h at 4 °C and stored at −80 °C until further use (20 mM HEPES pH 7.9, 0.2 mM EDTA, 0.5 mM DTT, 20 % glycerol with 250 mM KCl for Prp16 and 300 mM KCl for Prp22).

### Preparation of recombinant Prp18 protein in bacteria

The coding sequence for Prp18 was cloned into pET14. The protein was expressed as N-terminal His6-thrombin-Prp18 in *E. coli* BL21 (DE3)RIL cells. The protein was first purified on NiNTA-agarose beads and peak fractions dialyzed against 10 mM KPhosphate, 50 mM KCl pH 7.4. The protein was further purified on a hydroxyapatite column, and eluted with 0.5-3.0% ammonium sulfate. Finally, the purified protein was dialyzed into a suitable buffer for addition to splicing reactions (20 mM HEPES pH 7.9, 0.2 mM EDTA, 0.5 mM DTT, 20 % glycerol, 250 mM KCl).

### Splicing extract preparation and *in vitro* splicing

Yeast splicing extract was prepared using the liquid nitrogen method and *in vitro* splicing reactions were performed essentially as described (Lin et al., 1985; Umen & Guthrie, 1995). The resulting RNA species were phenol extracted and analyzed on 5 % (for *ACT1*) or 10 % (for *UBC4*) denaturing polyacrylamide gels. Quantification of gel bands was done using ImageJ (Rasband, 1997).

### Protein depletion from splicing extract

Prp18-3xHA was depleted from splicing extract by increasing the KCl concentration to 750 mM. The extract was incubated twice with 1/10 volume of anti-HA magnetic beads at 4 °C for several hours, followed by dialysis against 20 mM HEPES pH 7.9, 0.2 mM EDTA, 0.5 mM DTT, 20 % glycerol, 200 mM KCl for 16 hours. To ensure complete depletion of Prp18 while preventing co-depletion of Slu7, Western blots against Prp18-3xHA and Slu7-9xcmyc were performed. To rescue the splicing defect caused by Prp18-depletion, 60 ng of Prp18 were added to splicing reactions basically as described before (Horowitz & Krainer, 1997).

### Spliceosome assembly and purification

Complexes were assembled in a 1.5 mL splicing reaction on the indicated pre-mRNA substrate pre-bound to MS2-MBP fusion protein as described (Zhou et al., 2002). For reactions containing recombinantly expressed helicase mutants, the splicing extract was firstly incubated with 15 ng/mL final concentration of dominant negative mutant protein (Prp16-G378A or Prp22-K512A). Reactions were incubated at 23 °C for 30 min (*UBC4* transcripts) or 60 min (*ACT1* transcripts). *Complex C and C*:* The reaction mixture was adjusted to 2 mM glucose and incubation prolonged for 5 min. *Complex P:* To remove spliceosomes before the complex P stage, reactions were incubated with 5 uM of DNA complementary to the 3’-exon as described before (Wilkinson et al., 2017).

For all spliceosome preparations, the reaction mixture was centrifuged through a 40 % glycerol cushion in buffer A (20 mM HEPES, pH 7.9, 75 mM KCl, 0.25 mM EDTA). The cushion was collected and applied to amylose resin. After 15 hours of incubation at 4 °C, the resin was washed and eluted in 200 uL buffer A containing 5 % glycerol, 0.01 % NP-40 and 12 mM maltose. *Complex C*:* The beads were incubated with 2 mM ATP/2mM MgCl_2_ for 30 min at room temperature before washing (Fica et al., 2017).

### Spliceosome crosslinking, immunoprecipitation and cDNA library preparation

For each experiment, at least two replicates were produced from independently assembled and purified spliceosomes. In addition, control samples containing no tag on the protein and non-irradiated samples were prepared. Crosslinking and immunoprecipitation were adapted from the original iCLIP protocol (Huppertz et al., 2014) including the following modifications. 200 uL spliceosome elution were irradiated by UV light with a dose between 312 × 100 μJ/cm_2_ and 625 × 100 μJ/cm_2_ dependent on the protein of interest, the RNA transcript and the spliceosomal complex using a Stratalinker 2400 at 254 nm. 72 mg urea were added to denature the complex. The protein of interest was captured by magnetic anti-FLAG beads. The beads were incubated in 20 % S100 (a gift from S. Fica) in debranching buffer for 15 min to debranch intron-lariat as described (Ruskin & Green, 1985). 0.5 - 5 units (depending on complex and splicing transcript) of RNase I was added over 3 min at 37 °C for fragmentation. Dephosphorylation, adapter ligation, radioactive labeling, isolation of the RNA-protein complex and cDNA library preparation were performed as in iCLIP. cDNA libraries were sequenced at the Illumina HiSeq 2500 platform with cycle lengths between 100 and 250 nt in single-read mode.

### Data processing

A custom psiCLIP software pipeline was developed in Snakemake (Köster & Rahmann, 2012) and is available from github.com/luslab/psiclip. The code is easily extended to other experimental scenarios and substrates. Reads were demultiplexed using iCount demultiplex and trimmed for quality using Trim Galore! (Krueger, 2017). Quality of sequencing data was assessed with FastQC (Andrews & Others, 2010). Trimmed reads were mapped to a custom transcriptome index consisting of *S. cerevisiae* U1, U2, U4, U5 and U6 snRNAs alongside the pre-mRNA substrate sequence, using STAR aligner (Dobin et al., 2013). Key parameters were: `--alignEndsType EndToEnd` to prevent soft-clipping of reads which would obscure the truncation site, `--outFilterMismatchNmax 2` to allow a maximum of two mismatches and `--seedSearchStartLmax 16` which means the read is split into more seeds resulting in a potentially more sensitive search given our short reads. Subsequently, Sambamba was used to retrieve reads mapping only in the forward orientation to the custom transcriptome, as psiCLIP is stranded (Tarasov et al., 2015). Finally, the cDNA start position minus 1 is taken as the crosslink site and bed files are generated using Bedtools and R (Quinlan & Hall, 2010). Reads that did not map to the custom transcriptome index were then mapped to the yeast genome (SacCer3) and similarly processed to generate crosslinks. These crosslinks were used to normalise substrate and snRNA mapping crosslinks, as we found that they behaved as an internal spike in. Further plotting and analysis is performed in R using dplyr, ggplot2 and smoother packages (Hamilton, 2015; Wickham, 2016; Wickham et al., 2015). In the main figures we show the sum of all replicates, where available, with replicates shown separately in the supplementary material.

### Data and software availability

Scripts for data pre-processing and downstream processing are available from github.com/luslab/psiclip. Data is available to download from ArrayExpress at E-MTAB-8895.

## References

Andrews, S., & Others. (2010). FastQC: a quality control tool for high throughput sequence data.

Bai, R., Wan, R., Yan, C., Lei, J., & Shi, Y. (2018). Structures of the fully assembled Saccharomyces cerevisiae spliceosome before activation. Science, 360(6396), 1423–1429.

Blazquez, L., Emmett, W., Faraway, R., Pineda, J. M. B., Bajew, S., Gohr, A., Haberman, N., Sibley, C. R., Bradley, R. K., Irimia, M., & Ule, J. (2018). Exon Junction Complex Shapes the Transcriptome by Repressing Recursive Splicing. Molecular Cell, 72(3), 496–509.e9.

Briese, M., Haberman, N., Sibley, C. R., Faraway, R., Elser, A. S., Chakrabarti, A. M., Wang, Z., König, J., Perera, D., Wickramasinghe, V. O., Venkitaraman, A. R., Luscombe, N. M., Saieva, L., Pellizzoni, L., Smith, C. W. J., Curk, T., & Ule, J. (2019). A systems view of spliceosomal assembly and branchpoints with iCLIP. Nature Structural & Molecular Biology, 26(10), 930–940.

Burke, J. E., Longhurst, A. D., Merkurjev, D., Sales-Lee, J., Rao, B., Moresco, J. J., Yates, J. R., 3rd, Li, J. J., & Madhani, H. D. (2018). Spliceosome Profiling Visualizes Operations of a Dynamic RNP at Nucleotide Resolution. Cell, 173(4), 1014–1030.e17.

Chen, W., Moore, J., Ozadam, H., Shulha, H. P., Rhind, N., Weng, Z., & Moore, M. J. (2018). Transcriptome-wide Interrogation of the Functional Intronome by Spliceosome Profiling. Cell, 173(4), 1031–1044.e13.

Company, M., Arenas, J., & Abelson, J. (1991). Requirement of the RNA helicase-like protein PRP22 for release of messenger RNA from spliceosomes. Nature, 349(6309), 487–493.

Cordin, O., & Beggs, J. D. (2013). RNA helicases in splicing. RNA Biology, 10(1), 83–95.

Dobin, A., Davis, C. A., Schlesinger, F., Drenkow, J., Zaleski, C., Jha, S., Batut, P., Chaisson, M., & Gingeras, T. R. (2013). STAR: ultrafast universal RNA-seq aligner. Bioinformatics, 29(1), 15–21.

Fica, S. M., Oubridge, C., Galej, W. P., Wilkinson, M. E., Bai, X. C., Newman, A. J., & Nagai, K. (2017). Structure of a spliceosome remodelled for exon ligation. Nature, 542(7641), 377–380.

Galej, W. P., Wilkinson, M. E., Fica, S. M., Oubridge, C., Newman, A. J., & Nagai, K. (2016). Cryo-EM structure of the spliceosome immediately after branching. Nature, 537(7619), 197–201.

Giaever, G., Chu, A. M., Ni, L., Connelly, C., Riles, L., Véronneau, S., Dow, S., Lucau-Danila, A., Anderson, K., André, B., Arkin, A. P., Astromoff, A., El-Bakkoury, M., Bangham, R., Benito, R., Brachat, S., Campanaro, S., Curtiss, M., Davis, K.,…Johnston, M. (2002). Functional profiling of the Saccharomyces cerevisiae genome. Nature, 418(6896), 387–391.

Guthrie, C., & Patterson, B. (1988). Spliceosomal snRNAs. Annual Review of Genetics, 22, 387–419.

Haberman, N., Huppertz, I., Attig, J., König, J., Wang, Z., Hauer, C., Hentze, M. W., Kulozik, A. E., Le Hir, H., Curk, T., Sibley, C. R., Zarnack, K., & Ule, J. (2017). Insights into the design and interpretation of iCLIP experiments. Genome Biology, 18(1), 7.

Hamilton, N. (2015). smoother: Functions Relating to the Smoothing of Numerical Data. R Package Version 1.1. https://cran.rstudio.com/web/packages/smoother/index.html

He, Y., Staley, J. P., Andersen, G. R., & Nielsen, K. H. (2017). Structure of the DEAH/RHA ATPase Prp43p bound to RNA implicates a pair of hairpins and motif Va in translocation along RNA. RNA, 23(7), 1110–1124.

Horowitz, D. S., & Krainer, A. R. (1997). A human protein required for the second step of pre-mRNA splicing is functionally related to a yeast splicing factor. Genes & Development, 11(1), 139–151.

Huppertz, I., Attig, J., D’Ambrogio, A., Easton, L. E., Sibley, C. R., Sugimoto, Y., Tajnik, M., Konig, J., & Ule, J. (2014). iCLIP: protein-RNA interactions at nucleotide resolution. Methods, 65(3), 274–287.

James, S. A., Turner, W., & Schwer, B. (2002). How Slu7 and Prp18 cooperate in the second step of yeast pre-mRNA splicing. RNA, 8(8), 1068–1077.

Kondo, Y., Oubridge, C., van Roon, A.-M. M., & Nagai, K. (2015). Crystal structure of human U1 snRNP, a small nuclear ribonucleoprotein particle, reveals the mechanism of 5’ splice site recognition. eLife, 4. https://doi.org/10.7554/eLife.04986

Koodathingal, P., Novak, T., Piccirilli, J. A., & Staley, J. P. (2010). The DEAH box ATPases Prp16 and Prp43 cooperate to proofread 5’ splice site cleavage during pre-mRNA splicing. Molecular Cell, 39(3), 385–395.

Köster, J., & Rahmann, S. (2012). Snakemake—a scalable bioinformatics workflow engine. Bioinformatics, 28(19), 2520–2522.

Krueger, F. (2017). Trim Galore! http://www.bioinformatics.babraham.ac.uk/projects/trim_galore/

Kunkel, T. A. (1985). Rapid and efficient site-specific mutagenesis without phenotypic selection. Proceedings of the National Academy of Sciences of the United States of America, 82(2), 488–492.

Lee, F. C. Y., & Ule, J. (2018). Advances in CLIP Technologies for Studies of Protein-RNA Interactions. Molecular Cell, 69(3), 354–369.

Lee, S. C.-W., & Abdel-Wahab, O. (2016). Therapeutic targeting of splicing in cancer. Nature Medicine, 22(9), 976–986.

Leung, A. K. W., Nagai, K., & Li, J. (2011). Structure of the spliceosomal U4 snRNP core domain and its implication for snRNP biogenesis. Nature, 473(7348), 536–539.

Lin, R. J., Newman, A. J., Cheng, S. C., & Abelson, J. (1985). Yeast mRNA splicing in vitro. The Journal of Biological Chemistry, 260(27), 14780–14792.

Mayas, R. M., Maita, H., Semlow, D. R., & Staley, J. P. (2010). Spliceosome discards intermediates via the DEAH box ATPase Prp43p. Proceedings of the National Academy of Sciences of the United States of America, 107(22), 10020–10025.

McPheeters, D. S., & Muhlenkamp, P. (2003). Spatial organization of protein-RNA interactions in the branch site-3’ splice site region during pre-mRNA splicing in yeast. Molecular and Cellular Biology, 23(12), 4174–4186.

McPheeters, D. S., Schwer, B., & Muhlenkamp, P. (2000). Interaction of the yeast DExH-box RNA helicase Prp22p with the 3’ splice site during the second step of nuclear pre-mRNA splicing. Nucleic Acids Research, 28(6), 1313–1321.

Moore, M. J., & Sharp, P. A. (1992). Site-specific modification of pre-mRNA: the 2’-hydroxyl groups at the splice sites. Science, 256(5059), 992–997.

Ohrt, T., Odenwalder, P., Dannenberg, J., Prior, M., Warkocki, Z., Schmitzova, J., Karaduman, R., Gregor, I., Enderlein, J., Fabrizio, P., & Luhrmann, R. (2013). Molecular dissection of step 2 catalysis of yeast pre-mRNA splicing investigated in a purified system. RNA, 19(7), 902–915.

Plaschka, C., Newman, A. J., & Nagai, K. (2019). Structural Basis of Nuclear pre-mRNA Splicing: Lessons from Yeast. Cold Spring Harbor Perspectives in Biology, 11(5). https://doi.org/10.1101/cshperspect.a032391

Price, S. R., Evans, P. R., & Nagai, K. (1998). Crystal structure of the spliceosomal U2B“-U2A’ protein complex bound to a fragment of U2 small nuclear RNA. Nature, 394(6694), 645–650.

Query, C. C., & Konarska, M. M. (2004). Suppression of multiple substrate mutations by spliceosomal prp8 alleles suggests functional correlations with ribosomal ambiguity mutants. Molecular Cell, 14(3), 343–354.

Quinlan, A. R., & Hall, I. M. (2010). BEDTools: a flexible suite of utilities for comparing genomic features. Bioinformatics, 26(6), 841–842.

Rasband, W. S. (1997). ImageJ. http://www.worldlibrary.in/articles/eng/ImageJ

Rogelj, B., Easton, L. E., Bogu, G. K., Stanton, L. W., Rot, G., Curk, T., Zupan, B., Sugimoto, Y., Modic, M., Haberman, N., Tollervey, J., Fujii, R., Takumi, T., Shaw, C. E., & Ule, J. (2012). Widespread binding of FUS along nascent RNA regulates alternative splicing in the brain. Scientific Reports, 2, 603.

Ruskin, B., & Green, M. R. (1985). An RNA processing activity that debranches RNA lariats. Science, 229(4709), 135–140.

Schneider, S., Hotz, H.-R., & Schwer, B. (2002). Characterization of dominant-negative mutants of the DEAH-box splicing factors Prp22 and Prp16. The Journal of Biological Chemistry, 277(18), 15452–15458.

Schwer, B. (2008). A conformational rearrangement in the spliceosome sets the stage for Prp22-dependent mRNA release. Molecular Cell, 30(6), 743–754.

Schwer, B., & Gross, C. H. (1998). Prp22, a DExH-box RNA helicase, plays two distinct roles in yeast pre-mRNA splicing. The EMBO Journal, 17(7), 2086–2094.

Schwer, B., & Guthrie, C. (1992). A conformational rearrangement in the spliceosome is dependent on PRP16 and ATP hydrolysis. The EMBO Journal, 11(13), 5033–5039.

Semlow, D. R., Blanco, M. R., Walter, N. G., & Staley, J. P. (2016). Spliceosomal DEAH-Box ATPases Remodel Pre-mRNA to Activate Alternative Splice Sites. Cell, 164(5), 985–998.

Semlow, D. R., & Staley, J. P. (2012). Staying on message: ensuring fidelity in pre-mRNA splicing. Trends in Biochemical Sciences, 37(7), 263–273.

Sugimoto, Y., Konig, J., Hussain, S., Zupan, B., Curk, T., Frye, M., & Ule, J. (2012). Analysis of CLIP and iCLIP methods for nucleotide-resolution studies of protein-RNA interactions. Genome Biology, 13(8), R67.

Sutandy, F. X. R., Ebersberger, S., Huang, L., Busch, A., Bach, M., Kang, H.-S., Fallmann, J., Maticzka, D., Backofen, R., Stadler, P. F., Zarnack, K., Sattler, M., Legewie, S., & König, J. (2018). In vitro iCLIP-based modeling uncovers how the splicing factor U2AF2 relies on regulation by cofactors. Genome Research, 28(5), 699–713.

Tanaka, N., & Schwer, B. (2005). Characterization of the NTPase, RNA-binding, and RNA helicase activities of the DEAH-box splicing factor Prp22. Biochemistry, 44(28), 9795–9803.

Tarasov, A., Vilella, A. J., Cuppen, E., Nijman, I. J., & Prins, P. (2015). Sambamba: fast processing of NGS alignment formats. Bioinformatics, 31(12), 2032–2034.

Tseng, C.-K., & Cheng, S.-C. (2008). Both catalytic steps of nuclear pre-mRNA splicing are reversible. Science, 320(5884), 1782–1784.

Ule, J., & Blencowe, B. J. (2019). Alternative Splicing Regulatory Networks: Functions, Mechanisms, and Evolution. Molecular Cell, 76(2), 329–345.

Umen, J. G., & Guthrie, C. (1995). Prp16p, Slu7p, and Prp8p interact with the 3’ splice site in two distinct stages during the second catalytic step of pre-mRNA splicing. RNA, 1(6), 584–597.

Wagner, J. D., Jankowsky, E., Company, M., Pyle, A. M., & Abelson, J. N. (1998). The DEAH-box protein PRP22 is an ATPase that mediates ATP-dependent mRNA release from the spliceosome and unwinds RNA duplexes. The EMBO Journal, 17(10), 2926–2937.

Wickham, H. (2016). ggplot2: Elegant Graphics for Data Analysis. Springer.

Wickham, H., Francois, R., Henry, L., Müller, K., & Others. (2015). dplyr: A grammar of data manipulation. R Package Version 0. 4, 3.

Wilkinson, M. E., Fica, S. M., Galej, W. P., Norman, C. M., Newman, A. J., & Nagai, K. (2017). Postcatalytic spliceosome structure reveals mechanism of 3’-splice site selection. Science, 358(6368), 1283–1288.

Wilkinson, M. E., Lin, P. C., Plaschka, C., & Nagai, K. (2018). Cryo-EM Studies of Pre-mRNA Splicing: From Sample Preparation to Model Visualization. Annual Review of Biophysics, 47, 175–199.

Will, C. L., & Luhrmann, R. (2011). Spliceosome structure and function. Cold Spring Harbor Perspectives in Biology, 3(7). https://doi.org/10.1101/cshperspect.a003707

Yan, C., Wan, R., Bai, R., Huang, G., & Shi, Y. (2016). Structure of a yeast step II catalytically activated spliceosome. Science. https://doi.org/10.1126/science.aak9979

Zhou, Z., Sim, J., Griffith, J., & Reed, R. (2002). Purification and electron microscopic visualization of functional human spliceosomes. Proceedings of the National Academy of Sciences of the United States of America, 99(19), 12203–12207.

