## Supplementary material for "PsiCLIP reveals dynamic RNA binding by DEAH-box helicases before and after exon ligation"

### Supplementary Materials

**Figure S1: *In vitro* prepared splicing substrates used in psiCLIP.** All transcripts were generated to include 3 MS2 stem loops for spliceosome purification purposes. The transcripts are derived from the yeast *UBC4* (left) and *ACT1* gene (right). Distances between the recognition sequences are given in nt.

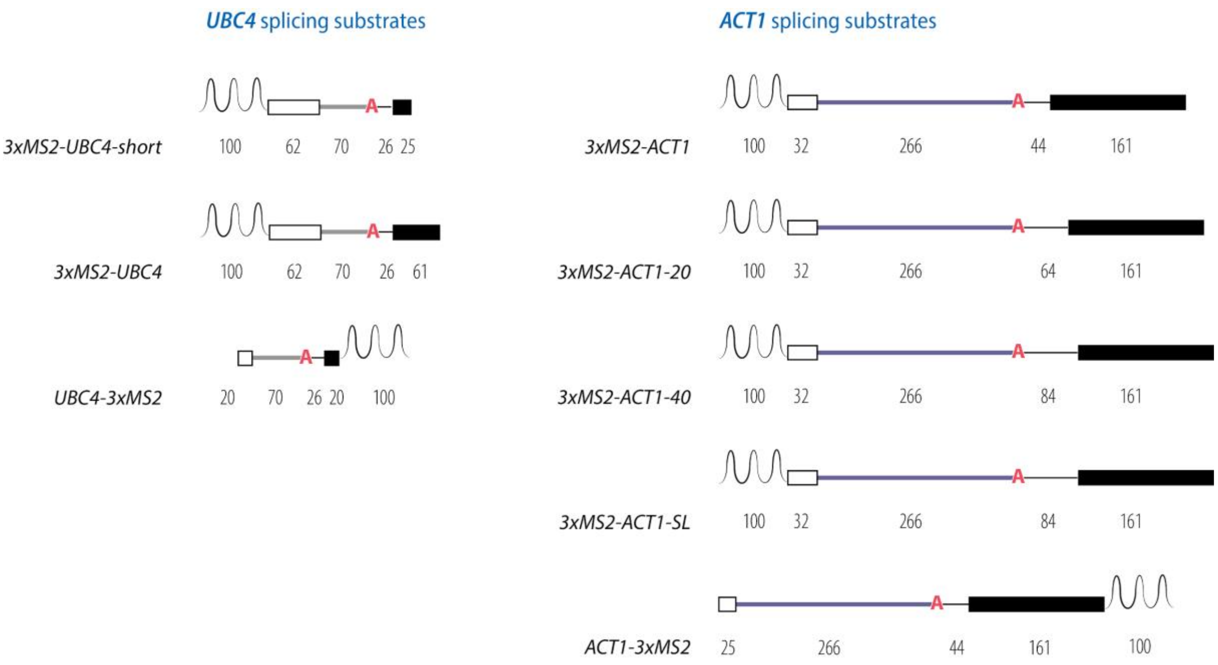

**Figure S2: DBR1 treatment resolves intron-lariats.** a) A schematic showing how intron-lariats can cause artefactual signal. If a protein (in blue) crosslinks near to the branch point, upon RNAse digestion part of the lariat may remain like a fork. In this situation, the free 3' end of the lariat is available for adaptor ligation (in orange). This means that in the final library there can be many reads that begin at the 5'SS and branchpoint that originate from the lariat and not a protein-RNA crosslink. b) Reverse transcription of the 5'SS guanosine and branch point adenosine can result in mutation. In the final reads, the branch point nucleotide is often read as a cytosine (top) and the 5'SS guanosine can be read as an adenosine or thymine (middle). For comparison, we show that the 3'SS adenosine is rarely mutated (bottom). DBR1 treatment reduces the incidence of mutations (right). Here we show 3 samples of DBR1- Prp16 psiCLIP, and 4 samples of DBR1+ Prp16 psiCLIP. c) Prp16 crosslinks in a DBR1- and DBR1+ sample, shown normalised to library size. The ratio between the artefact signal (branchpoint A in red) and true signal (a representative crosslink shown in blue) shifts dramatically upon DBR1 treatment.

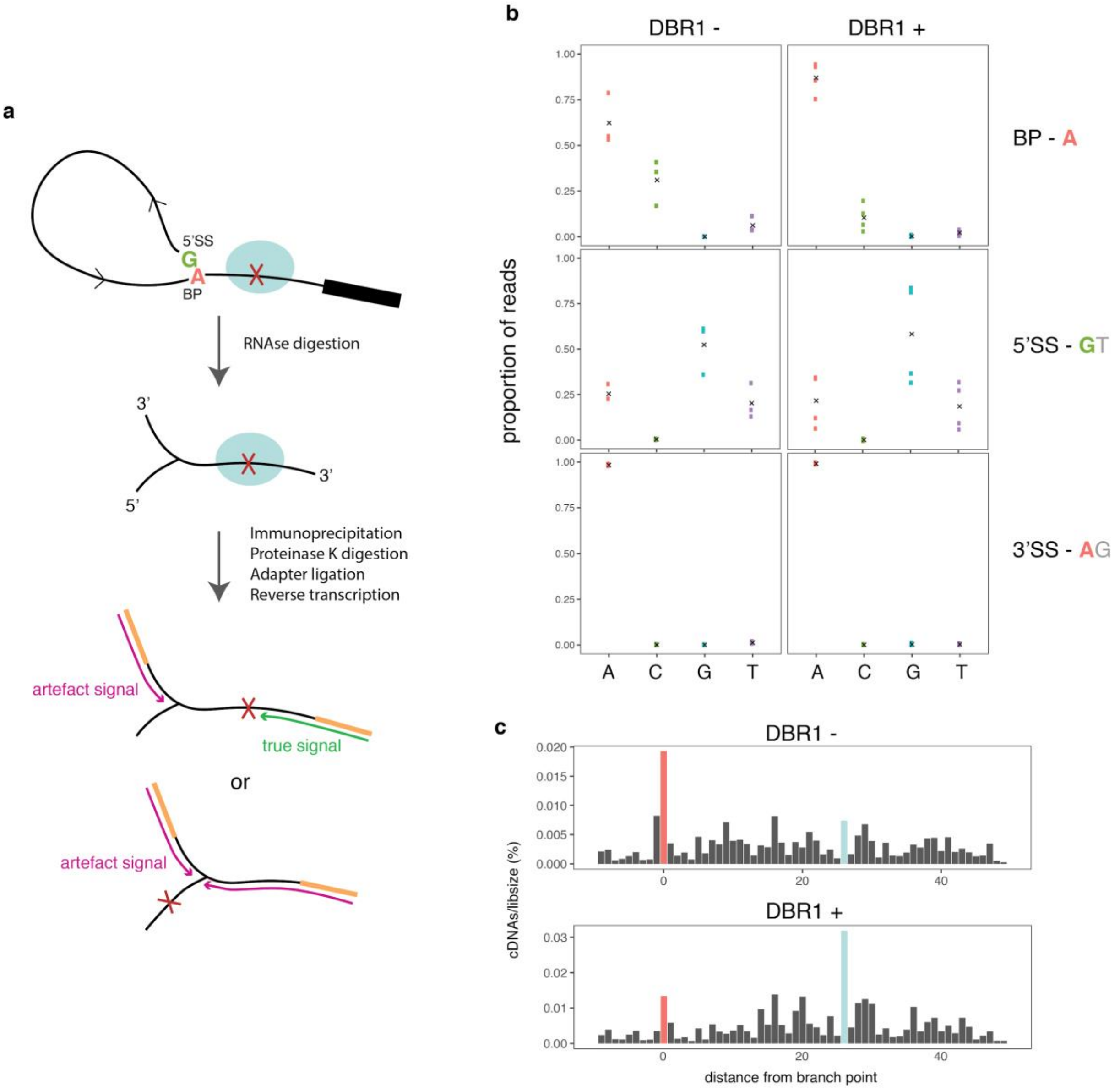

**Figure S3: Prp16 psiCLIP data show substrate specific binding in a region downstream of the brA**

**a)** Total number of crosslinks on the *ACT1* pre-mRNA substrate for three replicates of Prp16 psiCLIP. “AC” and “AG” denote mutant and canonical 3’SS respectively. **b)** Mapping of Prp16 psiCLIP data onto *ACT1*. Positions along the transcript are shown relative to the brA. The smoothed lines show the truncation events normalised to the number of cDNAs mapping to the yeast genome. We show an untagged control (ctrl, dotted lines) and the tagged experimental condition (UV, solid lines). Lines were gaussian smoothed with a window size of 20nt. **c)** Same as (a) but for three replicates of *UBC4* Prp16 psiCLIP. **d)** Same as (b) but for three replicates of *UBC4* Prp16 psiCLIP.

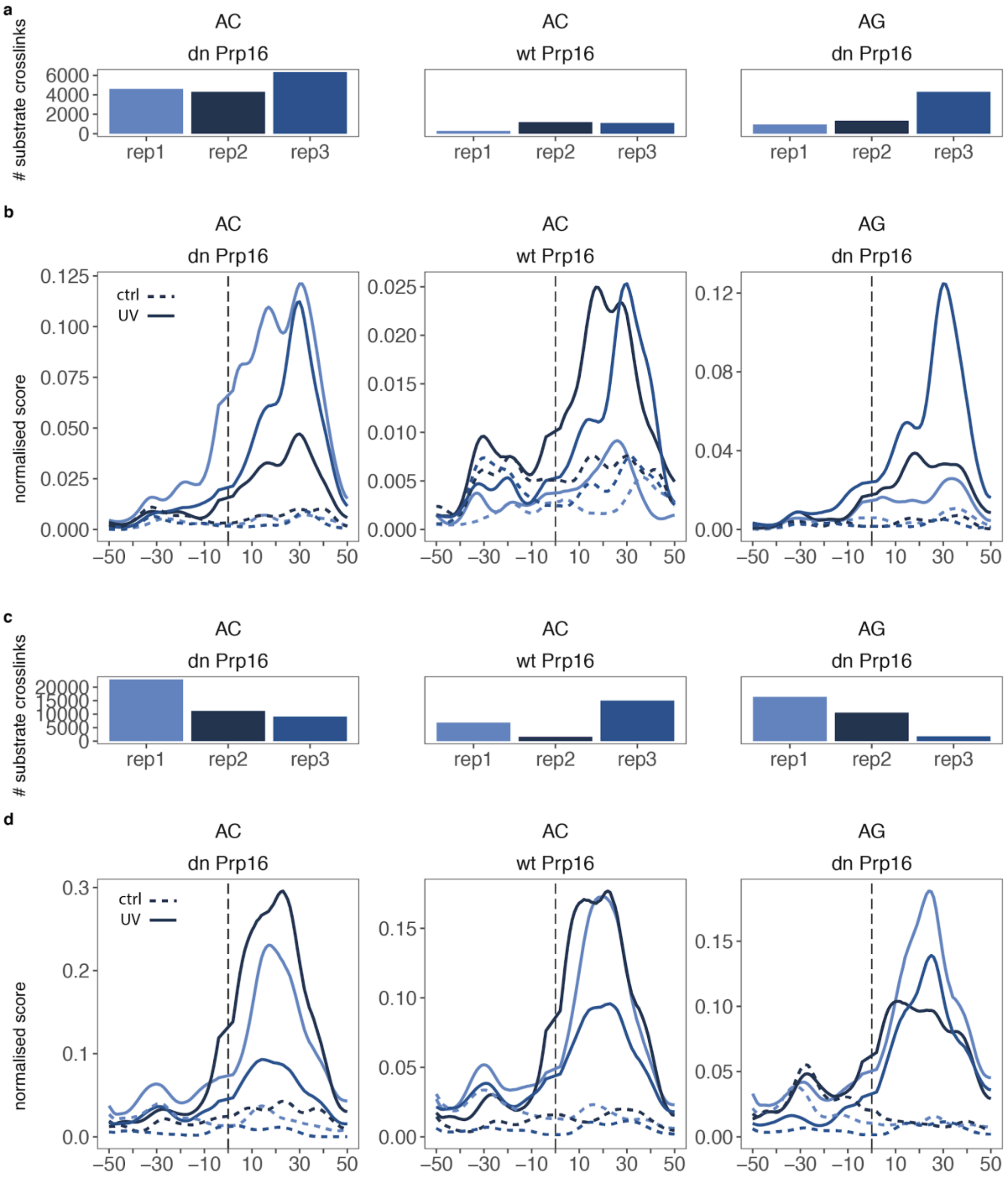

**Figure S4: Prp16 binds to accommodate expanded distance between BrA and 3'SS.** Two replicates for each condition are shown. Mapping of Prp16 mutant psiCLIP data onto the respective *ACT1* splicing substrate. cDNAs are normalised to total cDNAs mapping to the yeast genome. Crosslinks are shown with a 20nt window gaussian smooth. Positions on the substrate are given relative to the brA. Solid lines are tagged experimental conditions and dashed lines are untagged controls. **a)** Prp16-G378A on the WT *ACT1* substrate. **b)** Prp16-G378A on *ACT1* substrate with a 20nt extension between BrA and 3'SS. **c)** Prp16-G378A on *ACT1* substrate with a 40nt extension between BrA and 3'SS. **d)** Prp16-G378A on *ACT1* substrate with a stem loop in the 40nt extension between BrA and 3'SS.

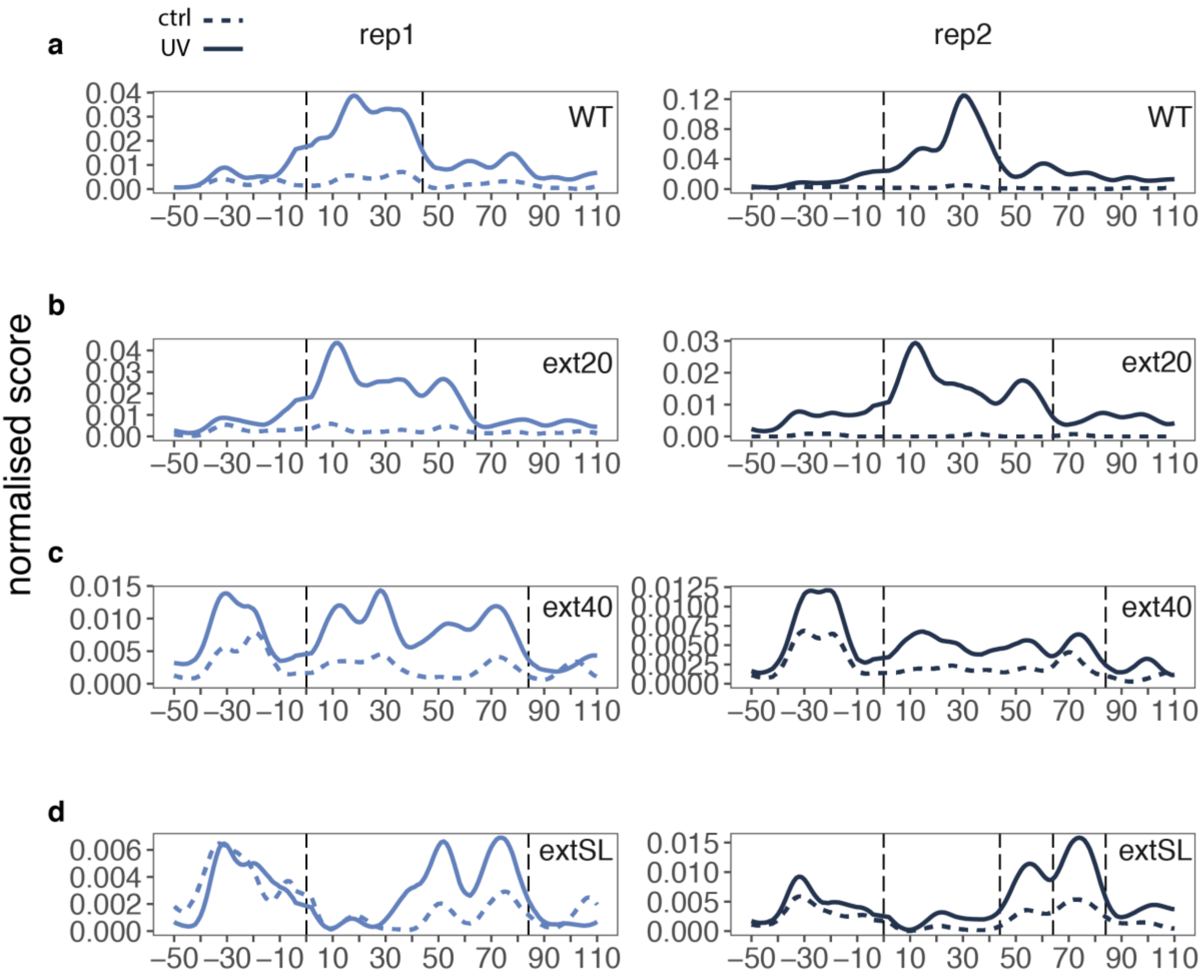

**Figure S5: Pattern of Prp22 psiCLIP data in C* and P complex is consistent between replicates. a)** Two replicates for each condition are shown. Mapping of Prp22 (WT and mutant) psiCLIP data onto the respective splicing substrate. cDNAs are normalised to total cDNAs mapping to the yeast genome. Crosslinks are shown with a 10nt window gaussian smooth. Positions on the substrate are given relative to the 3'SS. P complex crosslinks are shown with the intron removed. Solid lines are tagged experimental conditions and dashed lines are untagged controls. **b)** The proportion of spliced reads in each library showing that C* complex libraries have very few spliced reads, whereas P complex libraries are enriched for spliced reads. The colour denotes whether the splice junction is annotated - in other words the canonical splicing expected, or non-annotated. This shows that the spliceosome is using the canonical splice sites as expected in the P complex data.

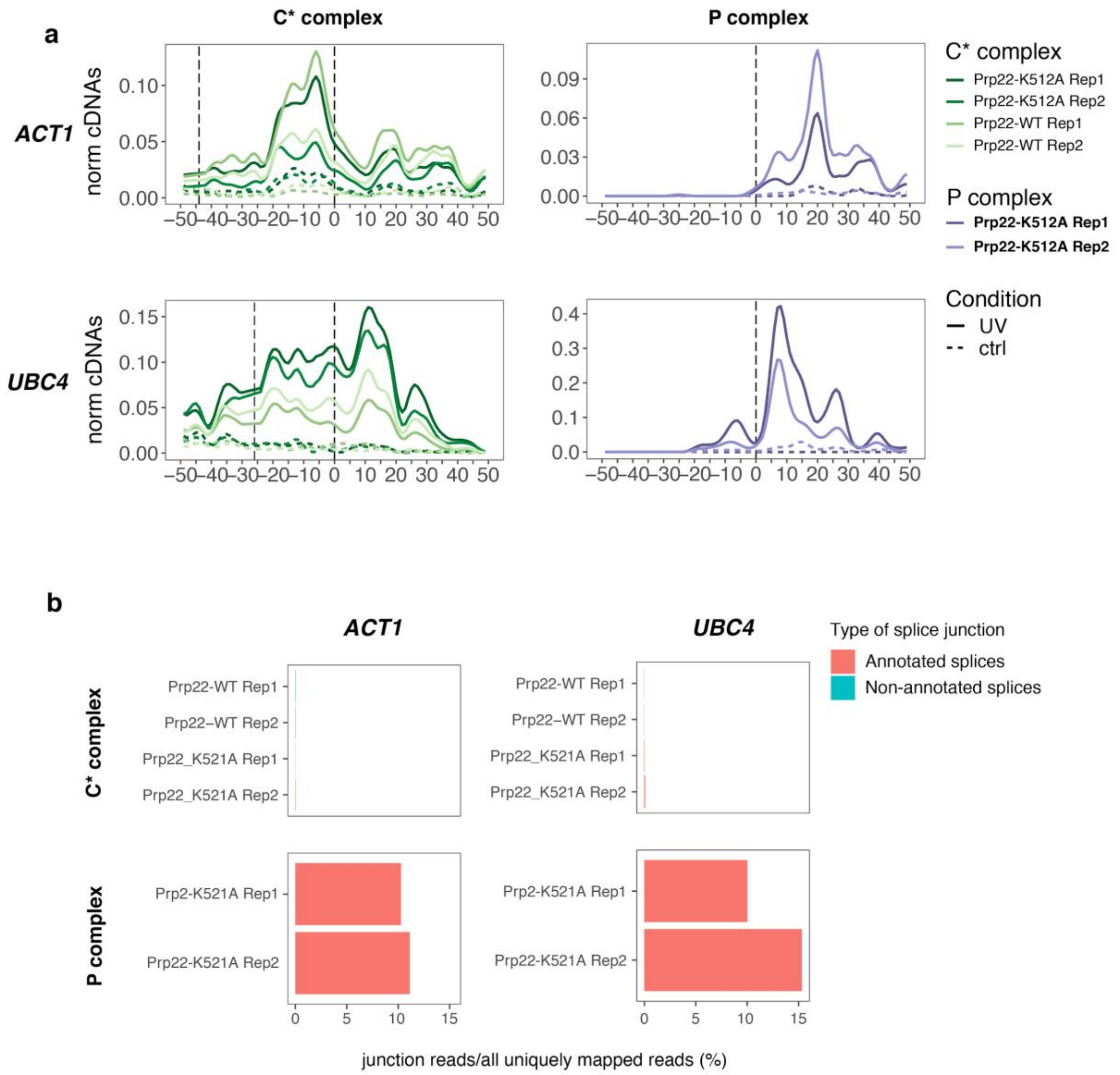

**Figure S6: Prp22 profile under Prp18 immunodepletion is consistent between replicates. a)** Denaturing gel after splicing assays with 3xMS2-*UBC4* demonstrate that there is a step two splicing defect under Prp18 depletion both in the background of Prp22-WT protein or with the addition of the dominant negative Prp22-K215A. The second step splicing defect can be rescued after 45 minutes by *in vitro* addition of recombinant Prp18 and is shown with the Prp22 WT background. **b)** dG substrate with Prp18 depletion. Denaturing gel of the spliceosomes eluted from amylose column showing the input for UV-crosslinking and immunoprecipitation. The amount of spliceosomes of the first step are reduced for Prp22 WT background compared to Prp22-K512A. **c)** Canonical AG substrate with Prp18 depletion. Denaturing gel of the spliceosomes eluted from amylose column showing the input for UV-crosslinking and immunoprecipitation. The amount of spliceosomes of the first step are reduced for Prp22 WT background compared to Prp22-K512A. **d-f)** Crosslinks are shown as normalised to the total yeast genome mapping reads and with the untagged ctrl subtracted from the tagged UV condition, with a 20nt window gaussian smooth. Replicate 1 data is shown on the left, and replicate 2 data is shown on the right. **d)** Prp22 psiCLIP in a WT genetic background on an *ACT1* substrate with dG mutation. Both WT and ATPase mutant Prp22 are shown. **e)** Prp22 psiCLIP under Prp18 depletion on an *ACT1* substrate with wildtype AG 3'SS. Both WT and ATPase mutant Prp22 are shown. **f)** Prp22 psiCLIP under Prp18 depletion on an *ACT1* substrate with dG mutation. Both WT and ATPase mutant Prp22 are shown. **g)** The proportion of spliced reads in each Prp18 depletion library showing that some splicing is able to proceed with the WT substrate, but that this is more enriched with the Prp22-K512A mutant. The colour denotes whether the splice junction is annotated - in other words the canonical splicing expected, or non-annotated. This shows that the majority of splicing we see is canonical.

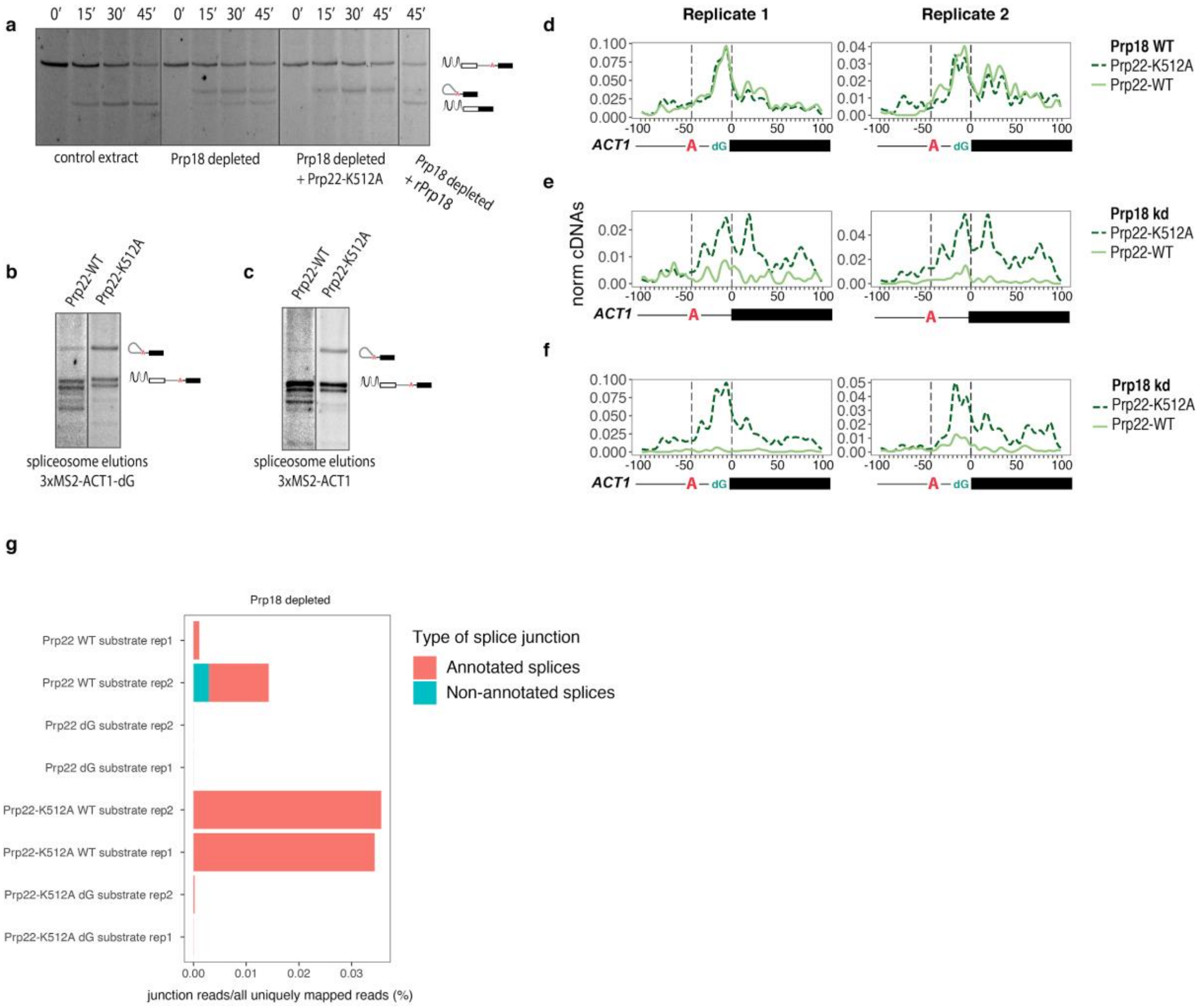
